# The Fanconi anemia pathway repairs colibactin-induced DNA interstrand cross-links

**DOI:** 10.1101/2024.01.30.576698

**Authors:** Maria Altshuller, Xu He, Elliot J. MacKrell, Kevin M. Wernke, Joel W. H. Wong, Selene Sellés-Baiget, Ting-Yu Wang, Tsui-Fen Chou, Julien P. Duxin, Emily P. Balskus, Seth B. Herzon, Daniel R. Semlow

## Abstract

Colibactin is a secondary metabolite produced by bacteria present in the human gut and is implicated in the progression of colorectal cancer and inflammatory bowel disease. This genotoxin alkylates deoxyadenosines on opposite strands of host cell DNA to produce DNA interstrand cross-links (ICLs) that block DNA replication. While cells have evolved multiple mechanisms to resolve (“unhook”) ICLs encountered by the replication machinery, little is known about which of these pathways promote resistance to colibactin-induced ICLs. Here, we use *Xenopus* egg extracts to investigate replication-coupled repair of plasmids engineered to contain site-specific colibactin-ICLs. We show that replication fork stalling at a colibactin-ICL leads to replisome disassembly and activation of the Fanconi anemia ICL repair pathway, which unhooks the colibactin-ICL through nucleolytic incisions. These incisions generate a DNA double-strand break intermediate in one sister chromatid, which can be repaired by homologous recombination, and a monoadduct (“ICL remnant”) in the other. Our data indicate that translesion synthesis past the colibactin-ICL remnant depends on Polη and a Polκ-REV1-Polζ polymerase complex. Although translesion synthesis past colibactin-induced DNA damage is frequently error-free, it can introduce T>N point mutations that partially recapitulate the mutation signature associated with colibactin exposure *in vivo*. Taken together, our work provides a biochemical framework for understanding how cells tolerate a naturally-occurring and clinically-relevant ICL.

## Introduction

Human health is profoundly influenced by host-microbiome interactions and the expansion of certain microbes is frequently linked to pathological states. Emerging data indicate many of these pathogenic gut bacteria contribute to dysbiosis and promote inflammation through the production of toxins that damage host DNA^1^. For example, pathogenic *pks^+^* bacteria harbor a 52 kilobase (kb) non-ribosomal polypeptide synthetase/polyketide synthase biosynthetic gene cluster that enables production of the colibactin genotoxin^2^. Colibactin-producing bacteria are detected in ∼40% of patients with inflammatory bowel disease (IBD) and ∼60% of patients with colorectal cancer and the overall abundance of these bacteria tends to be much higher in patients than in healthy subjects^3,4^. Moreover, colibactin exposure is associated with single-base substitution (SBS) and insertion/deletion (ID) mutation signatures found in the colonic crypts of most healthy individuals^5–7^. Colibactin is therefore a highly prevalent and potent driver of human disease.

*pks^+^* bacteria can produce multiple secondary metabolites that can alkylate DNA. However, the bioactive colibactin species (clb) that is thought to be primarily responsible for *pks^+^* bacteria pathogenicity comprises a pseudo-symmetric scaffold with two cyclopropane groups^8–10^. Each cyclopropane can react with a deoxyadenosine (dA) at the N3 position to form a DNA interstrand cross-link (ICL) that covalently links the two strands of DNA and blocks DNA unwinding. Consistently, clb-associated mutation signatures target AT-rich motifs containing offset dAs on opposite strands^6,7^. Once formed, the colibactin-ICL (clb-ICL) renders the alkylated dAs susceptible to depurination, resulting in the presence of N3 monoalkylated bases and DNA abasic (AP) sites that can further decompose into DNA strand breaks^11^. Thus, *pks^+^* bacteria likely generate a complicated spectrum of host DNA lesions that may mobilize multiple DNA repair pathways.

The clb-ICL is likely to be highly cytotoxic due to its presumed ability to stall replication forks. Failure to repair ICLs encountered during replication can lead to under-replication of the genome and/or gross chromosomal rearrangements. Consequently, cells have evolved multiple repair mechanisms that are activated when replication forks stall at an ICL (Figure 1A). An ICL induced by acetaldehyde (AA-ICL) can be resolved (“unhooked”) by an as yet unidentified enzyme that catalyzes direct reversal of the cross-link^12^. ICLs formed by psoralen and those formed by AP sites can be unhooked by the NEIL3 DNA glycosylase, which excises a cross-linked base to produce an AP site^13^. Should these pathways fail, the ICL is unhooked by the Fanconi anemia (FA) pathway, named for the bone marrow failure and cancer predisposition syndrome caused by mutations in this pathway^14^.

**Figure 1.**
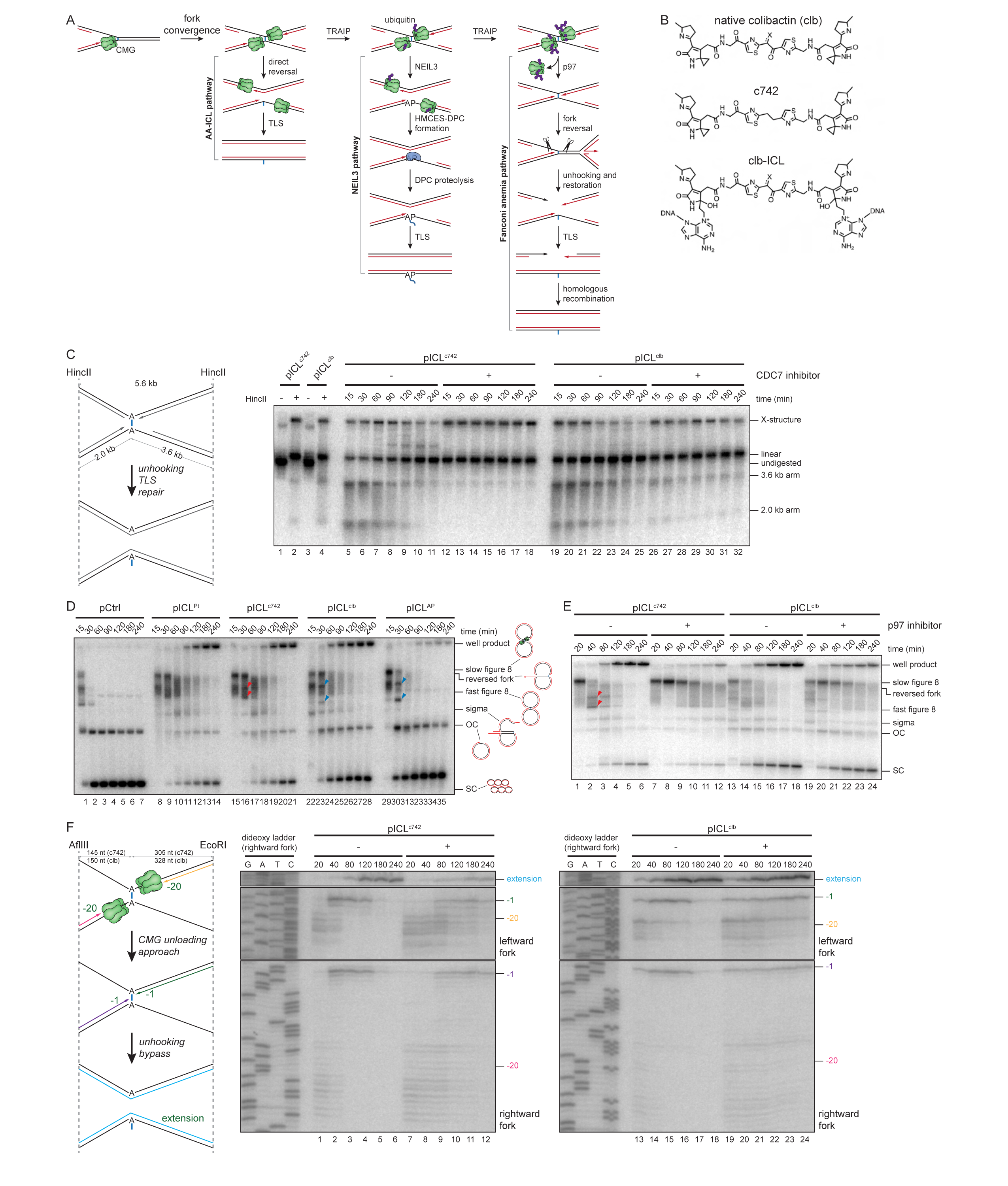
Replication-coupled repair of colibactin-induced ICLs. A. Schematic of replication-coupled ICL repair pathways (see main text for details). Black, parental DNA strands. Red, nascent DNA strands. Blue, ICL. Purple, ubiquitin. B. Structures of the native clb toxin (top; X indicates O or NH_2_), stabilized c742 colibactin analog (middle), and a potential structure of the clb-ICL. C. Left, schematic of species generated during ICL repair. Digestion with HincII produces X-structures corresponding to cross-linked plasmids and linear species corresponding to unhooked plasmids. Right, pICL^c742^ and pICL^clb^ were replicated in egg extract that was supplemented with CDC7 inhibitor, as indicated. Replication intermediates were recovered at the indicated times, digested with HincII and resolved by denaturing (alkaline) agarose gel electrophoresis. Intermediates were detected by Southern blotting with ^32^P radiolabeled probes and visualized by autoradiography. D. The indicated plasmids were replicated in egg extract supplemented with [α-^32^P]dCTP. Replication intermediates were separated on a native agarose gel and visualized by autoradiography. Schematics of replication and repair intermediates that are resolved by native agarose gel electrophoresis are shown. SC, supercoiled; OC, open circular. Red arrowheads indicate top and bottom intermediates formed during replication of pICL^c742^; blue arrowheads indicate OC-OC catenae and OC-SC catenae produced during replication of pICL^AP^ and pICL^clb^. E. pICL^c742^ and pICL^clb^ were replicated in egg extract supplemented with [α-^32^P]dCTP and p97i, as indicated. Replication intermediates were analyzed as in Figure 1D. Red arrowheads indicate top and bottom intermediates formed during replication of pICL^c742^. F. Left, schematic of nascent leading strands generated during ICL repair. AflIII cuts 145 or 150 nt to the left of the c742-ICL or clb-ICL, respectively. EcoRI cuts 305 or 328 nt to the right of the c742-ICL or clb-ICL, respectively). Digestion with AflIII and EcoRI generates characteristic –20 to –40 stall, –1 stall, and strand extension products. Right, nascent DNA strands from the pICL^c742^ and pICL^clb^ replication reactions shown in Figure 1E were isolated at the indicated times, digested with AflIII and EcoRI, and resolved by denaturing polyacrylamide gel electrophoresis. Top, middle, and bottom panels show sections of the same gels to visualize extension, leftward leading strands, and rightward leading strands, respectively.

The FA pathway is activated when replicative CDC45-MCM2-7-GINS (CMG) helicases converge and stall at an ICL^15^. CMGs are then polyubiquitylated by the replisome-associated TRAIP E3 ubiquitin ligase, leading to p97-dependent unloading of the CMGs and the reversal of one of the two converged forks^16–18^. At this stage, the FA core complex monoubiquitylates the FANCI-FANCD2 heterodimer, which becomes clamped onto DNA near the ICL^19,20^.

Monoubiquitylated FANCI-FANCD2 recruits structure-specific nucleases including XPF-ERCC1 that unhook the ICL through nucleolytic incisions flanking the ICL^21^. Following restoration of the reversed fork, one sister chromatid contains an ICL remnant that can be bypassed by the REV1-Polζ translesion synthesis (TLS) polymerase while the other chromatid contains a DNA double strand break (DSB) that is repaired by homologous recombination^22,23^. Although the FA pathway is more complicated than the AA-ICL and NEIL3 repair pathways, it is thought to be capable of unhooking any ICL, regardless of the chemistry underlying cross-link formation, and is therefore regarded as the most important pathway for cellular resistance to ICL-inducing agents.

Despite the versatility of the FA pathway, the identities of physiological ICLs targeted by this pathway remain obscure. Genetic evidence indicates that deficits in aldehyde metabolism synergize with mutations in the FA pathway, implying that the FA pathway resolves an ICL formed by endogenous reactive aldehydes, such as formaldehyde or acetaldehyde^24,25^. Indeed, chemically-synthesized acetaldehyde-ICLs that escape unhooking by direct reversal are processed by the FA pathway in *Xenopus* egg extracts^12^. However, the formation of ICLs induced by acetaldehyde or any other endogenous metabolite has not yet been confirmed *in vivo*. Compounding the ambiguity surrounding the function of the FA pathway, FANC proteins have been shown to promote a variety of genome maintenance mechanisms beyond ICL repair, including homologous recombination, replication fork protection, R-loop resolution, retrotransposon suppression, and mitochondrial genome maintenance^26–30^. Intriguingly, indirect evidence suggests that clb-ICLs might be repaired by the FA pathway. Colibactin exposure induces FANCD2 ubiquitylation and FANCD2 foci formation in cells and FANCD2-deficient cells are hypersensitive to colibactin^31^. However, given the spectrum of DNA lesions produced during colibactin exposure and the diverse functions of the FA pathway, it remains unclear whether clb-ICLs are repaired by the FA pathway or whether the FA pathway influences the progression of colibactin-associated pathologies.

Here, we used *Xenopus* egg extracts to investigate replication-coupled repair of site-specific ICLs formed by either the native colibactin toxin or by a stabilized colibactin analog. We find that both ICLs are strong impediments to DNA replication and are repaired in a replication-coupled manner. Unhooking of the ICLs occurs through nucleolytic incisions mediated by the FA pathway. TLS past both an unhooked colibactin-induced ICL remnant and a colibactin monoadduct depends on REV1 and Polκ. We further show that colibactin-induced ICL repair can be error-free, but frequently introduces primarily T>A point mutations at the site of alkylation. These results elucidate the mechanisms of repair for DNA lesions that may drive the progression of IBD and CRC.

## Results

### Colibactin-ICL repair is coupled to DNA replication

To investigate colibactin-ICL repair in *Xenopus* egg extracts, we prepared plasmids containing site-specific ICLs generated with either native colibactin (clb) or a synthetic colibactin analog (c742) (Figure 1B). c742 contains an unsubstituted carbon-carbon linker in place of a central diketone that renders the native clb genotoxin susceptible to oxidative cleavage. The stabilized analog produces an analog that is otherwise identical to ICL formed by clb and can induce cell cycle arrest in human cells^32^. Short oligonucleotide duplexes containing a colibactin alkylation hotspot motif were incubated with either the c742 analog or with cultured *pks^+^* bacteria (Supplementary Figure 1, A and B). In both cases, cross-links were expected to form between dAs at the second position of the 5’-A**A**TATT-3’ palindromic hotspot sequence^6,7^. Purified cross-linked duplexes were then ligated into ∼5.6 kb plasmids, and the covalently-closed circular products were isolated on CsCl gradients. Analysis of the plasmids revealed that >75% of the c742-ICL plasmid (pICL^c742^) contained an intact ICL while >60% of the clb-ICL plasmid (pICL^clb^) contained an ICL (Supplementary Figure 1C). Interestingly, the non-cross-linked fraction of pICL^clb^ contained AP sites and monoadducts, presumably consisting of clb linked to adenine (clb-A) (Supplementary Figure 1, D and E). These observations indicate that the clb-ICL can undergo depurination during plasmid preparation. Consistent with previous work, the AP sites present in our pICL^clb^ preparations are rapidly removed in egg extracts (Supplementary Figure 1F)^33^. We therefore infer that replication forks encounter either a clb-ICL or clb-A monoadduct, but not an AP site, during pICL^clb^ replication in egg extract.

We first asked whether colibactin-induced ICLs are unhooked in egg extracts and whether any observed unhooking depends on DNA replication. pICL^c742^ and pICL^clb^ were replicated in egg extract plasmids, recovered, linearized with a restriction enzyme (HincII), and resolved on a denaturing agarose gel. The plasmid DNA was then detected by Southern blotting (Figure 1C). Initially, the plasmids were detected as “X-structures” that migrate slowly due to the presence of a cross-link. The initial accumulation of clb-ICL X-structures was reduced due to the presence of uncross-linked plasmid in our preparations (Supplementary Figure 1, C-E). During replication in egg extract, the X-structures are resolved into a linear species, indicating that the ICL in the DNA template is unhooked (Figure 1C, lanes 5-11 and 19-25). Importantly, the conversion of X-structures was attenuated in extract supplemented with CDC7 inhibitor (PHA-767491), which blocks the initiation of DNA replication (Figure 1C, lanes 12-18 and 26-32)^34^. We conclude that c742-ICL and clb-ICL are unhooked by replication-coupled repair in egg extracts.

To gain insights into the mechanism of colibactin-induced ICL repair, pICL^c742^ and pICL^clb^ were replicated in egg extract supplemented with [α-^32^P]dCTP (which is incorporated into the nascent DNA strands), and replication intermediates were resolved on a native agarose gel and visualized by autoradiography (Figure 1D). For comparison, we also replicated an undamaged plasmid (pCtrl) and plasmids containing either a cisplatin-ICL (pICL^Pt^) or an AP-ICL (pICL^AP^). Whereas replicated pCtrl rapidly accumulated as open circular and supercoiled products, each of the ICL-containing plasmids initially accumulated as slowly migrating “figure 8” intermediates that form when CMG helicases translocating along opposite strands converge and stall at the ICL. In the case of pICL^AP^, the slow figure 8s are rapidly converted into open circular and supercoiled products, indicative of unhooking by the NEIL3 glycosylase^13^. In the case of pICL^c742^ and pICL^clb^, the slow figure 8s were first converted into a series of species with intermediate mobilities and then into open circular and supercoiled products at later timepoints. Note that pICL^clb^ replication resulted in substantial accumulation of open circular and supercoiled plasmids at even the earliest time points, consistent with the presence of uncross-linked plasmid in this preparation. Interestingly, pICL^c742^ and pICL^clb^ replication produced intermediates that closely resembled those formed during replication of pICL^Pt^, which is known to be repaired by the FA pathway in egg extracts^35,36^. These findings indicate that colibactin-induced ICLs block replication fork progression and subsequently initiate repair.

We noticed that replication of pICL^c742^, but not pICL^clb^, consistently produced two prominent intermediates (top and bottom) that persisted throughout the replication time course. Our characterizations of these intermediates indicate that the two species accumulate after CMG unloading and approach of the nascent leading strands up to the ICL (Supplementary Figure 1G). We suspect that the novel intermediates correspond to topological isomers of cross-linked fast figure 8 structures that have not undergone fork reversal. We hypothesize that the structure of the c742-ICL might antagonize fork reversal or that the c742-ICL might be bound by proteins that impede fork reversal. Additional studies will be required to determine why this fraction of the c742-ICLs apparently fail to efficiently engage ICL repair machinery after CMG unloading.

In egg extracts, replication-coupled ICL repair requires the convergence of replication forks on either side of the ICL, which leads to TRAIP-dependent polyubiquitylation of CMG^16^. Short ubiquitin chains on CMG are sufficient to recruit the NEIL3 glycosylase, while longer chains trigger p97-dependent CMG unloading and activation of the FA pathway. We confirmed that colibactin-induced ICL repair requires fork convergence (Supplementary Figure 1H) and so tested whether it also requires CMG unloading. Addition of p97 inhibitor (NMS-873^37^) to pICL^c742^ and pICL^clb^ replication reactions caused persistent accumulation of slow figure 8s and a reduction in unhooked open-circular and supercoiled products (Figure 1E), indicating that repair is activated by CMG unloading. Importantly, the absence of unhooking in p97-inhibited extracts implies that the c742-and clb-ICLs are not processed by NEIL3 or AA-ICL pathways, whose activations do not require CMG unloading^12,13^. We also examined the effect of p97 inhibition on maturation of the nascent DNA strands by digesting repair intermediates with restriction enzymes (AflIII and EcoRI) that enable resolution of the leading strands on a denaturing sequencing gel (Figure 1F). During pICL^c742^ and pICL^clb^ replication, the nascent leading strands initially stall ∼20 to 40 nucleotides (nt) upstream of the ICL due to the footprint of CMG. The nascent strands are subsequently extended to 1 upstream of the expected dA cross-link positions before eventually being converted into full length extension products. Consistent with a requirement for CMG unloading during pICL^c742^ and pICL^clb^ repair, p97 inhibitor blocked extension of the nascent strands from –20 to –40 stall positions to the –1 position and suppressed the formation of full-length extension products. We conclude that colibactin-induced ICL repair is activated by fork convergence and CMG unloading, suggestive of repair by the FA pathway.

### Colibactin-induced ICLs are unhooked by the FA pathway

To determine which repair networks are activated during colibactin-induced ICL repair, we performed unbiased quantitative mass spectrometry on chromatin recovered from pICL^c742^ replication reactions (Figure 2A and Supplementary Figure 2, A and B)^38^. As expected, the CMG helicase and DNA synthesis machinery (e.g. Polδ and Polε polymerases) were enriched on chromatin at early, but not later, replication timepoints (Figure 2A and Supplementary Figure 2B), consistent with replisome disassembly during replication-coupled ICL repair. Dissociation of the replisome coincided with recruitment of proteins implicated in the FA pathway, including ATR-ATRIP, FA core complex proteins, FANCI-FANCD2, structure-specific nucleases, TLS polymerases, and homologous recombination proteins. Importantly, FANCI-FANCD2, structure-specific nucleases, TLS polymerases, and homologous recombination proteins did not accumulate on pCtrl or on pICL^c742^ that was incubated in non-replicating egg extracts (Figure 2A). Consistently, we also observed that replication of pICL^c742^ and pICL^clb^, but not pCtrl, stimulated mono-ubiquitylation of FANCD2 in egg extracts (Supplementary Figure 2C). For reasons we do not understand, FANCM and the FA core complex appeared to be more abundant on both pCtrl and non-replicating pICL^c742^. However, this may reflect an artifact of our analysis, since FANCM and the FA core complex increase in abundance during pICL^c742^ replication in a manner that temporally correlates with replisome unloading and the association of other FA pathway proteins. Taken together, these data indicate that the FA pathway is mobilized during replication-coupled repair of pICL^c742^.

**Figure 2.**
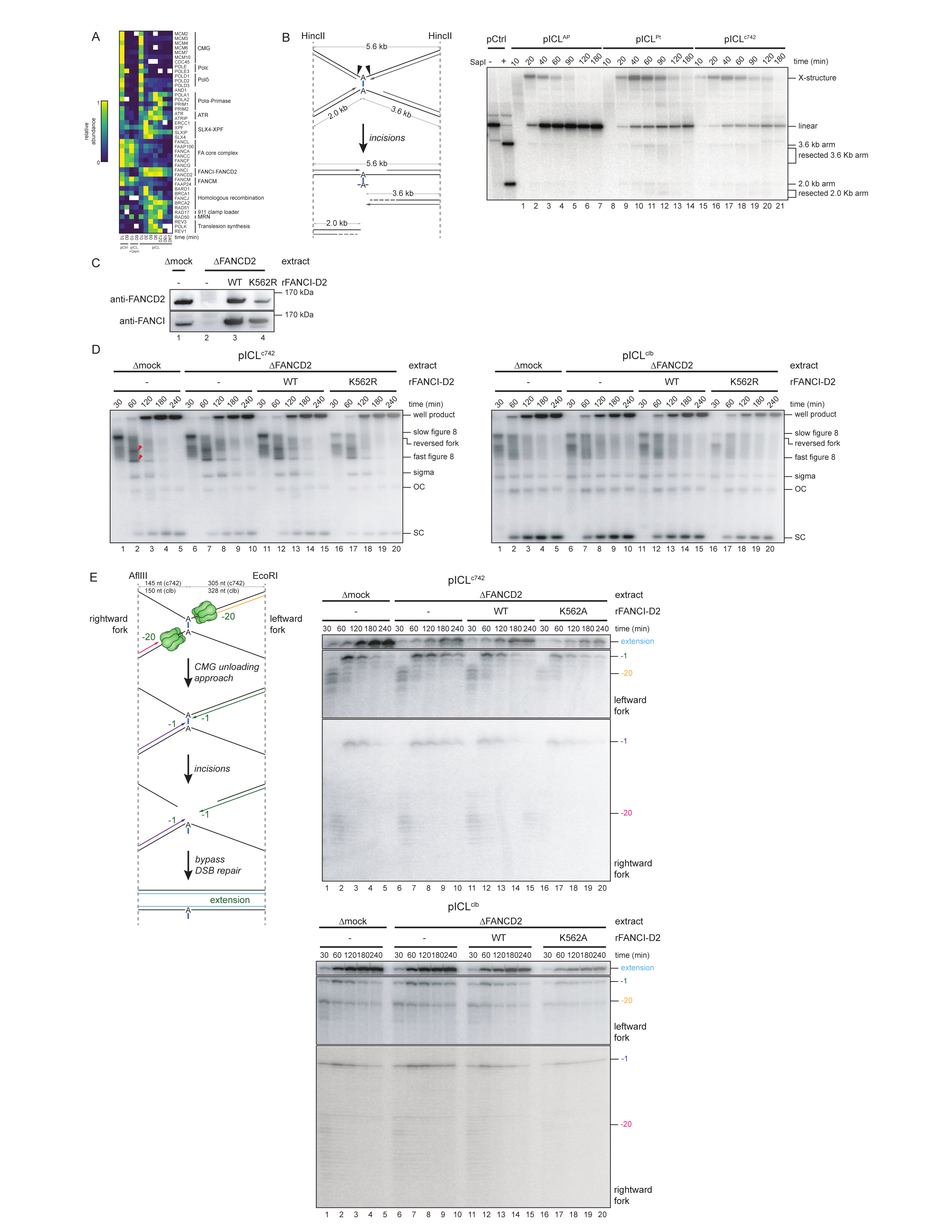
FA pathway activation during colibactin-induced ICL repair. A. The indicated plasmids were incubated in egg extract supplemented with Geminin, as indicated. At various timepoints, the plasmids were recovered from extract (in triplicate) and chromatin-associated proteins were quantified by mass spectrometry. The heat map indicates the relative abundance of proteins following global normalization. White boxes indicate that the protein was not detected. B. Left, schematic of incision products generated during nucleolytic ICL repair. Digestion with HincII produces X-structures corresponding to cross-linked plasmids and linear species corresponding to unhooked plasmids. Nucleolytic incisions flanking the ICL produce ∼2.0 and ∼3.6 double strand brake fragments that are resolved on a native agarose gel. Right, pCtrl, pICL^c742^, and pICL^Pt^ were replicated in egg extract supplemented with [α-^32^P]dCTP. Replication intermediates were recovered at the indicated times, digested with HincII, resolved by native agarose gel electrophoresis, and visualized by autoradiography. C. FANCI-FANCD2 immunodepletion. The extracts used in the replication reactions shown in Figure 2,D and E were blotted for FANCI and FANCD2. D. pICL^c742^ and pICL^clb^ were replicated with [α-^32^P]dCTP in mock– or FANCI-FANCD2-depleted extracts supplemented with rFANCI-FANCD2, as indicated. Replication intermediates were analyzed as in Figure 1D. SC, supercoiled; OC, open circular. Red arrowheads indicate top and bottom intermediates formed during replication of pICL^c742^. E. Left, schematic of nascent leading strands generated during ICL repair by the FA pathway. Digestion with AflIII and EcoRI generates characteristic –20 to –40 stall, –1 stall, and strand extension products, as in Figure 1F. Right, nascent DNA strands from the pICL^c742^ and pICL^clb^ replication reactions shown in Figure 2D were isolated at the indicated times, digested with AflIII and EcoRI, and analyzed as in Figure 1F.

ICL repair by the FA pathway is distinguished by nucleolytic incisions that effectively convert the ICL into a DSB. To test whether replication-coupled repair of colibactin-induced ICLs involves nucleolytic unhooking, we replicated pICL^c742^ in egg extracts supplemented with [α-^32^P]dCTP. Recovered replication and repair intermediates were linearized and then resolved on a native agarose gel (Figure 2B). Replicated pICL^c742^ initially accumulated as cross-linked X-structures that were converted into ∼2.0 and ∼3.6 kb fragments, consistent with DSB formation during nucleolytic repair. Over time, these fragments disappeared due to either repair by homologous recombination or nucleolytic degradation. The pattern of pICL^c742^ incision products accumulation was indistinguishable from the pattern of incision products generated during replication of pICL^Pt^, which is unhooked by the FA pathway. By contrast, incision products were not observed to appreciably accumulate during replication of pICL^AP^, which is predominantly unhooked without incisions by NEIL3. These data imply that colibactin-induced ICLs are unhooked through nucleolytic incisions mediated by the FA pathway. The resulting DSB is then presumably repaired by homologous recombination factors, which our data indicate are recruited to plasmids containing a colibactin-induced ICL (Figure 2A).

To explicitly test whether unhooking of the colibactin-induced ICLs requires FANCI-FANCD2, we replicated pICL^c742^ and pICL^clb^ in mock– or FANCI-FANCD2-immunodepleted extracts supplemented with [α-^32^P]dCTP and resolved replication and repair intermediates on a native agarose gel (Figure 2, C and D). FANCI-FANCD2-depletion resulted in a subtle but reproducible accumulation of reversed fork intermediates and a reduction in the amount of fully replicated supercoiled plasmid, indicating a defect in ICL repair by the FA pathway. Consistently, this effect was largely reversed by the addition of wild-type recombinant (r)FANCI-FANCD2 complex, but not by rFANCDI-FANCD2^K562R^, which cannot undergo FANCD2 mono-ubiquitylation and does not support nucleolytic incisions^21^. Analysis of the nascent DNA leading strands revealed FANCI-FANCD2-depletion caused a persistence of –1 stall products and a reduction of full-length products that was rescued by addition of wild-type rFANCI-FANCD2^WT^, but not by addition of rFANCI-FANCD2^K562R^ (Figure 2E). As in the case of a cisplatin-ICL (Supplementary Figure 2, D to F)^36^, FANCI-FANCD2 therefore enables extension of nascent leading strands beyond the colibactin cross-linked bases. Taken together, our data indicate that, following CMG unloading and fork reversal, FANCI-FANCD2 promotes nucleolytic incisions that unhook the cross-linked parental DNA strands, thereby enabling TLS past the resulting ICL remnant and completion of DNA replication.

### REV1 and Polη promote translesion synthesis during colibactin-induced ICL repair

Following unhooking of an ICL, TLS extends the nascent DNA strands past damaged nucleotides in the template DNA strands. In the case of ICL remnants formed by unhooking of nitrogen mustard– and cisplatin-ICLs, TLS is mediated by an insertion polymerase, which inserts a nucleotide opposite the damaged base, and the REV1-Polζ complex, which extends the DNA strand^22^. In contrast, both the insertion and extension steps require REV1 during TLS past ICL remnants formed by unhooking of acetaldehyde-ICLs as well as during bypass of AP sites and psoralen adducts produced by NEIL3-dependent unhooking^12,13^. REV1 immunodepletion largely abolished the formation of supercoiled plasmids during pICL^c742^ replication (Figure 3, A and B). This was accompanied by an accumulation of open circular plasmids, consistent with a requirement for REV1 during TLS past the unhooked c742-ICL. REV1 depletion also caused a pronounced accumulation of open circular plasmid during pICL^clb^ replication, indicating that bypass of either an unhooked clb-ICL remnant or the clb-A monoadduct also depends on REV1 (see below). Note that supercoiled plasmid still accumulated during pICL^clb^ replication in REV1-depleted extract, likely due the presence of undamaged template strands generated upon repair of AP sites opposite clb-A monoadducts. For both pICL^c742^ and pICL^clb^, REV1 depletion caused a strong persistence of –1 stall products and a reduction in full-length extension products (Figure 3C). This was distinct from the persistence of insertion (0 stall position) products observed during cisplatin-ICL repair in REV1-depleted extracts (Supplementary Figure 3, A-C)^22^. Our data therefore indicate that REV1 is required for both the insertion and extension TLS steps during repair of colibactin-induced ICLs. In some experiments, small amounts of insertion products were detected during bypass of the c742-ICL that were suppressed by REV1-depletion. These insertion products were not detected during bypass of the clb-ICL, suggesting that subtle differences in the structures of the c742– and clb-ICLs may influence the likelihood of REV1-dependent insertion opposite an alkylated dA comprising a colibactin-induced ICL.

**Figure 3.**
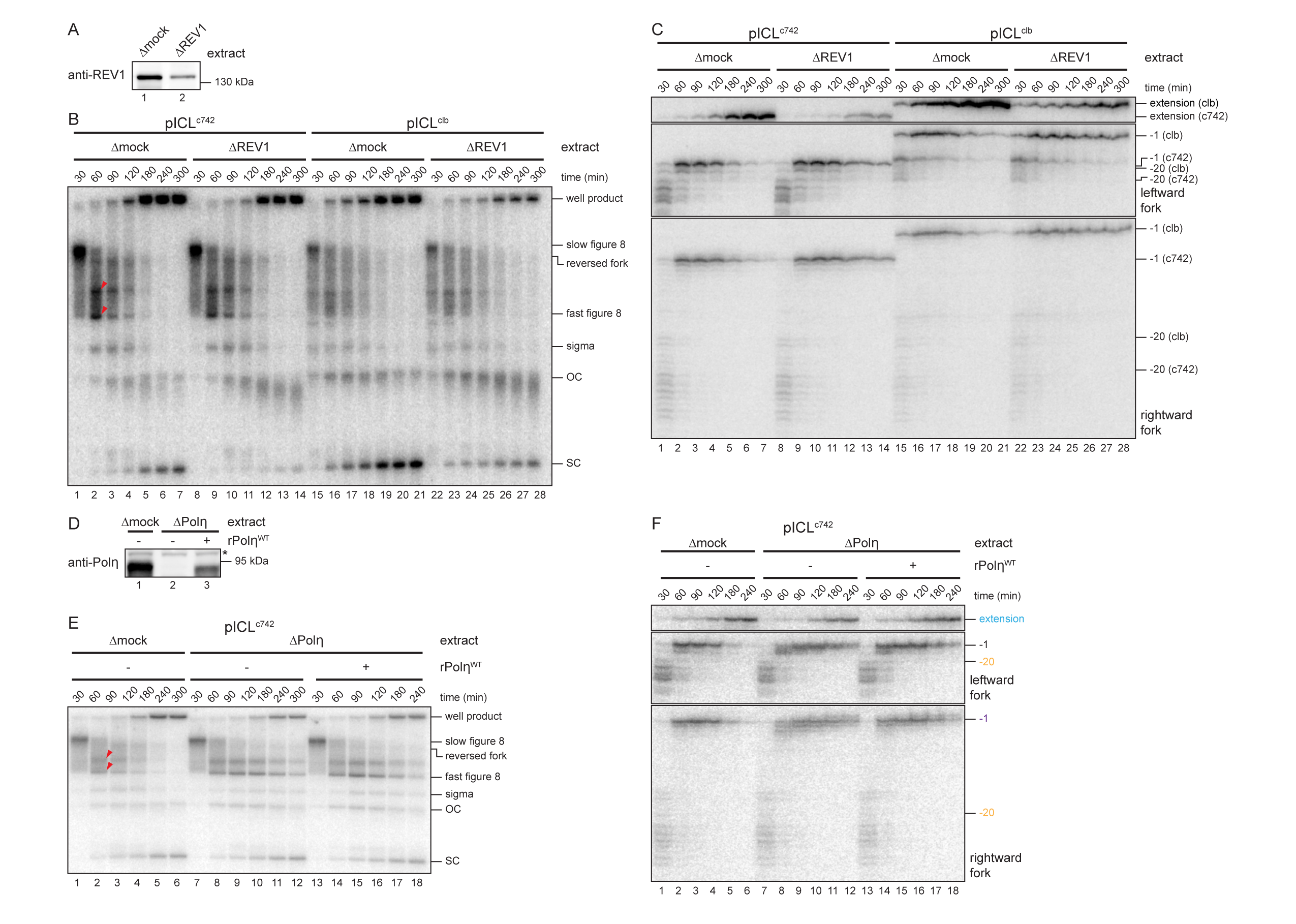
Translesion synthesis during colibactin-induced ICL repair. A. REV1 immunodepletion. The extracts used in the replication reactions shown in Figure 3, B and C were blotted for REV1. B. pICL^c742^ and pICL^clb^ were replicated with [α-^32^P]dCTP in mock– or REV1-depleted extracts and replication intermediates were analyzed as in Figure 1D. SC, supercoiled; OC, open circular. Red arrowheads indicate top and bottom intermediates formed during replication of pICL^c742^. C. Nascent DNA strands from the pICL^c742^ and pICL^clb^ replication reactions shown in 3B were isolated at the indicated times, digested with AflIII and EcoRI, and analyzed as in Figure 1F. D. Polη immunodepletion. The extracts used in the replication reactions shown in Figure 3, E and F were blotted for Polη. Asterisk indicates non-specific band. E. pICL^c742^ was replicated with [α-^32^P]dCTP in mock– or Polη –depleted extracts supplemented with rPolη, as indicated. Replication intermediates were analyzed as in Figure 1D. SC, supercoiled; OC, open circular. Red arrowheads indicate top and bottom intermediates formed during replication of pICL^c742^. F. Nascent DNA strands from the pICL^c742^ replication reactions shown in Figure 3E were isolated at the indicated times, digested with AflIII and EcoRI, and analyzed as in Figure 1F.

We also tested whether TLS during colibactin-induced ICL repair depends on Polη, which has been reported to function as an insertion polymerase during TLS past ICLs and other DNA lesions^39^. Polη immunodepletion did not appreciably affect the formation of supercoiled plasmids during pICL^c742^ replication in egg extracts (Figure 3,D and E), suggesting that Polη is dispensable for TLS during colibactin-induced ICL repair. However, Polη depletion did cause a transient accumulation of stall products from the –2 to the –7 position that was largely suppressed by addition of wild-type rPolη (Figure 3F). This pattern of stall product accumulation was similar to that observed during replication of pICL^Pt^ (Supplementary Figure 3, A-C). We conclude that Polη normally extends nascent leading strands to within 1 nt of a colibactin-induced ICL, but when Polη is absent, another DNA polymerase, such as Polδ, can perform this function, albeit less efficiently.

### TLS past a colibactin-induced monoadduct depends on REV1 and Pol**κ**

During preparation of pICL^clb^, depurination of the clb-ICL resulted in the formation of site-specific clb-A monoadducts and AP sites (that are rapidly removed in egg extract; Supplementary Figure 1, D to F). We exploited this decomposition of the clb-ICL to examine the mechanism of TLS past a colibactin-induced monoadduct. Previous work has implicated the polymerase activity of Polκ in TLS past minor groove adducts, including those formed by N3 dA alkylation^40,41^. We replicated pICL^clb^ in mock– or Polκ-depleted egg extract that was supplemented with p97 inhibitor to block CMG unloading and suppress approach of the nascent leading strands up to the ICL (Figure 4A). Under these conditions, TLS is expected to predominantly reflect encounters with clb-A monoadducts produced by depurination. Polκ depletion inhibited the formation of supercoiled plasmids during pICL^clb^ replication and caused an accumulation of nascent leading strand stall products at –1 relative to the expected positions of the clb-A monoadducts with a concomitant reduction in full length extension products (Figure 4, B and C). We therefore conclude that bypass of a clb-A adduct depends on Polκ.

**Figure 4.**
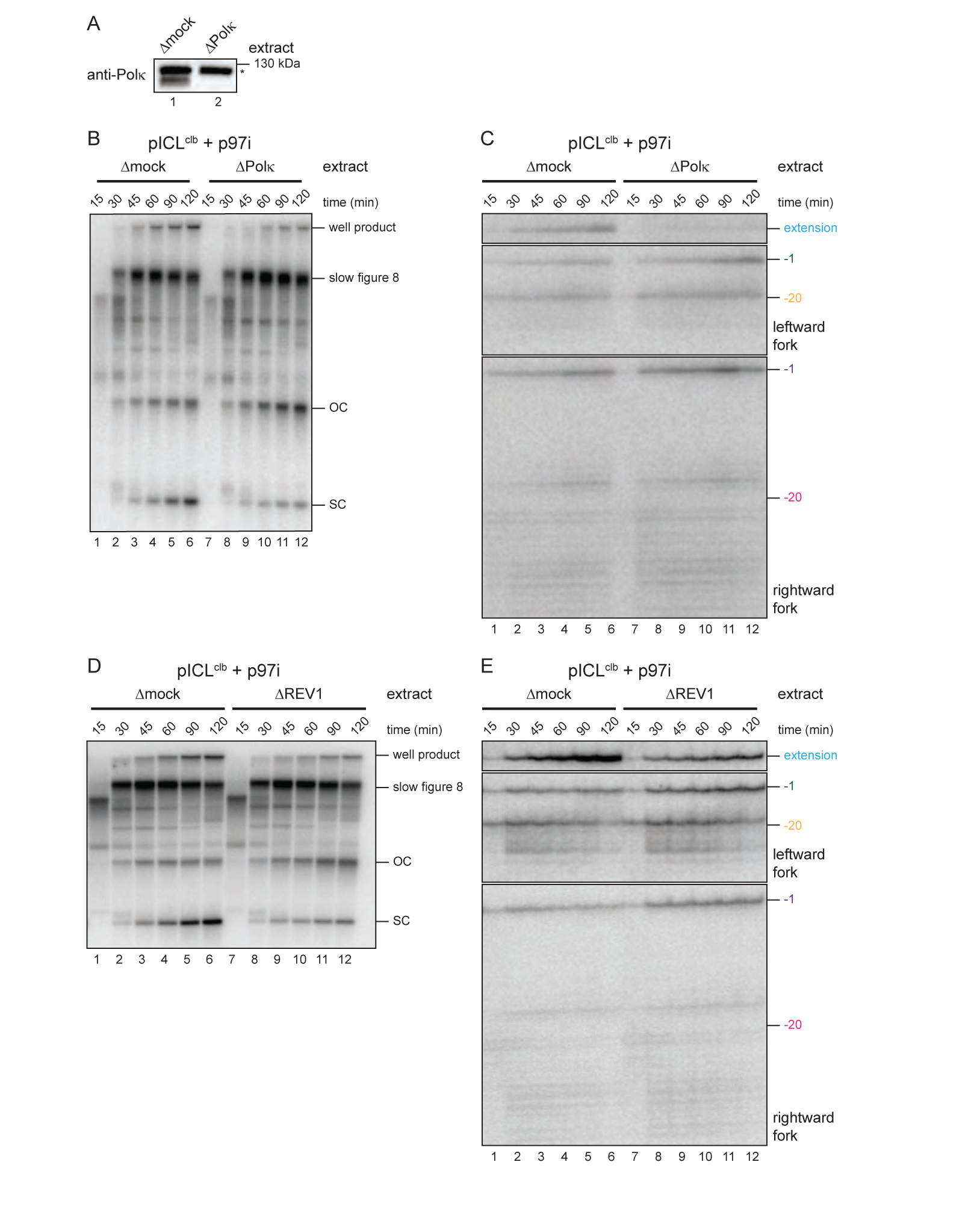
Translesion synthesis during bypass of a colibactin-induced monoadduct. A. Polκ immunodepletion. The extracts used in the replication reactions shown in Figure 4, B and C and Supplementary Figure 4B were blotted for Polκ and REV1. Asterisk indicates non-specific band. B. pICL^clb^ was replicated with [α-^32^P]dCTP in mock– or Polκ-depleted extracts that were supplemented with p97 inhibitor and replication intermediates were analyzed as in Figure 1D. SC, supercoiled; OC, open circular. C. Nascent DNA strands from the pICL^clb^ replication reactions shown in Figure 4B were isolated at the indicated times, digested with AflIII and EcoRI, and analyzed as in Figure 1F. D. pICL^clb^ was replicated with [α-^32^P]dCTP in mock– or REV1-depleted extracts that were supplemented with p97 inhibitor and replication intermediates were analyzed as in Figure 1D. SC, supercoiled; OC, open circular. E. Nascent DNA strands from the pICL^clb^ replication reactions shown in D were isolated at the indicated times, digested with AflIII and EcoRI, and analyzed as in Figure 1F.

Polκ acts independently of REV1 in egg extract to bypass previously examined minor groove adducts such as 3-deaza-3-phenethyl-adenosine acylfulvene (p3d-Phen-A)^41^. Polκ also functions non-catalytically to promote REV1-Polζ-dependent TLS past major groove adducts. We sought to test whether TLS past a clb-A adduct reflects a catalytic or non-catalytic requirement for Polκ. Unfortunately, technical issues have prevented us from testing whether the effects of Polκ-depletion are rescued by the addition of wild-type or catalytically-dead rPolκ proteins. Instead, we replicated pICL^clb^ in mock– or REV1-depleted egg extract supplemented with p97 inhibitor to test whether TLS past a clb-A monoadduct requires REV1 (Supplementary Figure 4A), which would be most consistent with a non-catalytic role for Polκ. Similar to Polκ-depletion, REV1-depletion also inhibited the formation of supercoiled plasmids and caused an accumulation of nascent leading strand stall products at –1 relative to the expected positions of the clb-A monoadducts with a concomitant reduction in full length extension products (Figure 4, D and E). These data imply that Polκ acts non-catalytically and in concert with REV1-Polζ to enable TLS past a clb-A monoadduct. We also observed that replication of pICL^c742^ in Polκ-depleted extract resulted in an accumulation of gapped, open-circular plasmids and a reduction in fully replicated, supercoiled plasmids (Supplementary Figure 4B), suggesting that unhooked c742-ICL remnants and clb-A are bypassed through a common REV1– and Polκ-dependent mechanism.

### Bypass of colibactin-induced DNA damage is mutagenic

Exposure to *pks^+^* bacteria produces T>N SBSs and 1 nt T deletions at inferred colibactin alkylation hotspots^5–7^, but the repair processes responsible for these mutation signatures have not been established. To determine whether colibactin-induced ICL repair by the FA pathway is mutagenic, we replicated pICL^c742^ in egg extract and performed next generation sequencing on recovered repair products (Figure 5A and Supplementary Figure 5, A and B). We also sequenced repair products recovered from replication reactions that were supplemented with p97 inhibitor to inactivate the FA pathway. Under unperturbed conditions, we observed that ∼60% of the recovered, sequenceable plasmids were free of mutations, insertions, and deletions, while 25% of plasmids contained point mutations within the AATATT colibactin hotspot (Figure 5B, left). The vast majority (∼90%) of these point mutations mapped to the expected alkylation sites, indicating that bypass of the unhooked colibactin-ICL remnant can be mutagenic. In cells and organoids, colibactin exposure is associated primarily with T>C SBSs, and less frequently with T>A and T>G mutations. Interestingly, c742-ICL repair produced mostly T>A mutations, followed by T>G, and T>C mutations. This suggests that while the FA pathway may account for colibactin-induced T>A mutations, another mutagenic process may generate the T>C mutations most associated with colibactin exposure. Addition of p97 inhibitor reduced the amount of fully repaired plasmid recovered from replication reactions but did not alter the overall frequency or spectrum of observed point mutations (Figure 5B, right). Residual activation of the FA pathway may therefore account for most single base substitutions generated in p97-inhibited extract.

**Figure 5.**
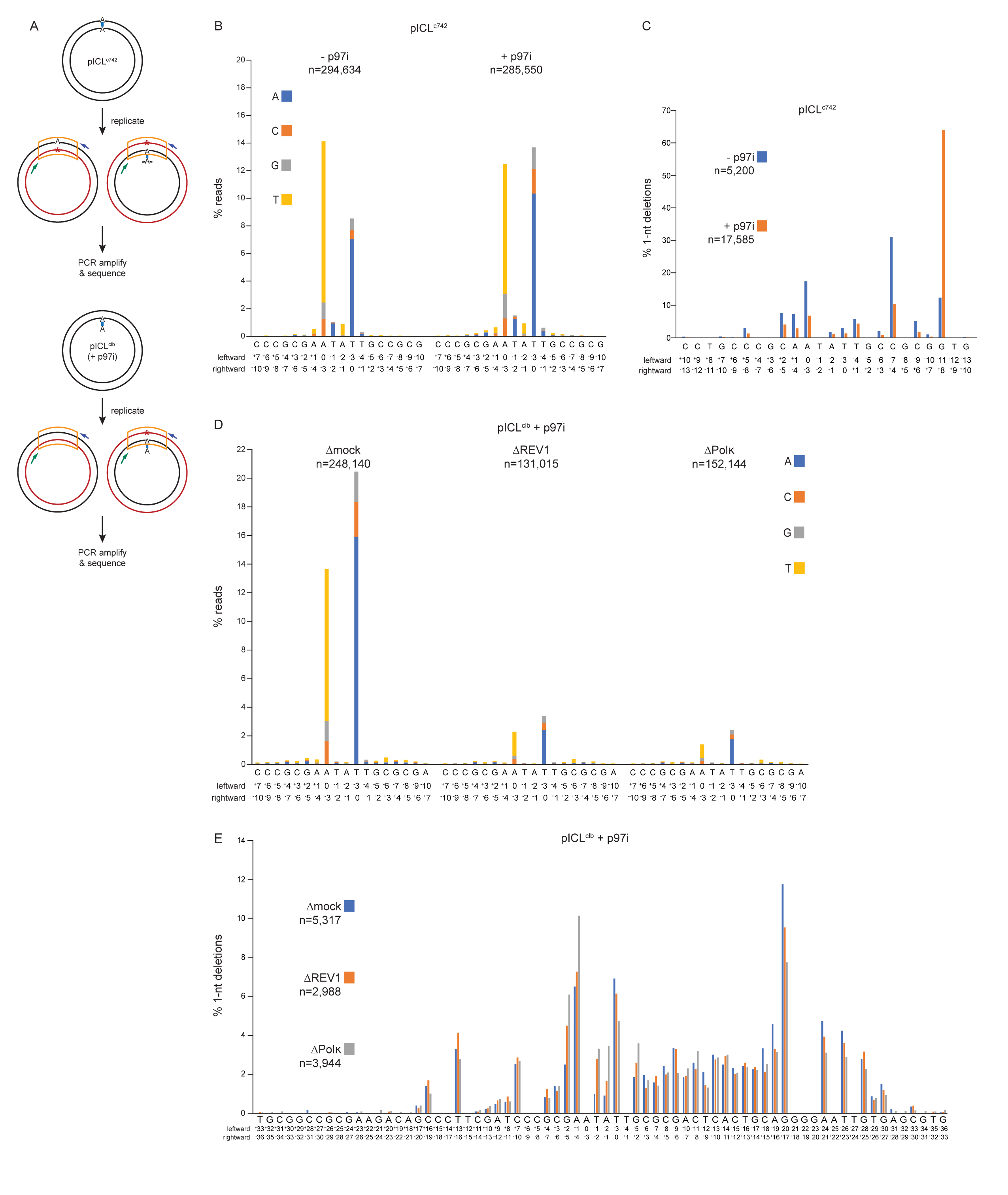
Mutagenicity of colibactin-induced DNA damage. A. Schematic depicting analysis of pICL^c742^ (top) and pICL^clb^ (bottom) repair products by next generation sequencing. Blue and green arrows indicate PCR primers; red asterisk indicates potential point mutation. B. pICL^c742^ was replicated in extract supplemented with p97 inhibitor, as indicated, and repair products were PCR amplified and sequenced. The fraction of reads corresponding to single base substitutions at each position in the vicinity of the ICL is plotted for each extract condition. n indicates the number of mapped sequencing reads obtained for each condition. The reference sequence is shown with numbers indicating the positions for the rightward and leftward nascent leading strands relative to the site of cross-linking (“0”). C. pICL^c742^ was replicated in extract supplemented with p97 inhibitor, as indicated, and repair products were PCR amplified and sequenced, as in Figure 5B. The fraction of single nucleotide deletions at each position in the vicinity of the ICL is plotted for each extract condition. n indicates the number of mapped reads containing a single nucleotide deletion. D. pICL^clb^ was replicated in the indicated extracts supplemented with p97 inhibitor and repair products were PCR amplified and sequenced. The fraction of reads corresponding to single base substitutions at each position in the vicinity of the ICL is plotted for each extract condition. n indicates the number of mapped sequencing reads obtained for each condition. The reference sequence is shown with numbers indicating the positions for the rightward and leftward nascent leading strands relative to the site of cross-linking (“0”). E. pICL^clb^ was replicated in the indicated extracts supplemented with p97 inhibitor and repair products were PCR amplified and sequenced, as in Figure 5D. The fraction of fraction of single nucleotide deletions at each position in the vicinity of the ICL is plotted for each extract condition. n indicates the number of mapped reads containing a single nucleotide deletion.

Deletion mutations were evident in ∼5% of pICL^c742^ repair products under unperturbed conditions. These deletions spanned 1 to 48 nt in length with a median length of 11 nt (Supplementary Figure 5C; –p97i condition). Single nucleotide deletions were the most common type of deletion (accounting for ∼33% all deletions) and clustered within ∼10 nt of the alkylated positions (Figure 5C and Supplementary Figure 5C; –p97i condition). Interestingly, blocking activation of the FA pathway with p97 inhibitor resulted in an overall increase in the frequency of deletion mutations, but did not substantially influence the distribution of deletion lengths (Supplementary Figure 5, C and D; compare –p97i and +p97i conditions). p97 inhibition also did not appreciably alter the distribution of single nucleotide deletions, which remained clustered within ∼10 nt the colibactin alkylation hotspot (Figure 5C). This finding implies that the observed single nucleotide deletions are not generated through FA pathway-dependent ICL repair. Combined, these data suggest that in the absence of CMG unloading and repair by the FA pathway, aberrant processing of stalled forks can introduce larger deletions.

Since the vast majority of fully replicated pICL^clb^ plasmids were generated upon replication of contaminating uncross-linked plasmids containing clb-A monoadducts (Figure 4), we were unable to distinguish the mutation spectrum produced during clb-ICL repair. However, this instead allowed us to examine the mutagenicity of the colibactin-induced monoadduct. We replicated pICL^clb^ in mock-, REV1-, or Polκ-depleted egg extract supplemented with p97 inhibitor (again, to inactivate ICL repair by the FA pathway) and sequenced the replication products (Supplementary Figure 5, E-G). Replication past clb-A monoadducts in mock-depleted extract produced a mutagenic tract that mirrored the tract produced by c742-ICL repair. We observed predominantly T>A mutations at the colibactin hotspot (Figure 5D). The frequency of single-base substitutions within the AATATT hotspot (35%) was also similar to that generated during c742-ICL repair, with 97% of these point mutations mapping to the expected dA alkylation sites. Depletion of either REV1 or Polκ dramatically reduced the frequency of hotspot point mutations (6% and 4% for REV1– and Polκ-depletion, respectively), indicating that REV1– and Polκ-dependent TLS is responsible for the observed spectrum of point mutations (Figure 5D). Note that in the absence of REV1 or Polκ, a greater proportion of sequencing reads are likely derived from undamaged strands present following AP site repair, resulting in an apparent reduction in the frequency of single-base substitutions during pICL^clb^ replication.

Bypass of the clb-A lesion in mock-depleted extract also introduced more broadly distributed deletions near the colibactin hotspot. These deletions ranged from 1-70 nt in length with single nucleotide deletions being most abundant (Supplementary Figure 5, H and I). Strikingly, the frequency of deletions rose from ∼4% in mock-depleted extract to ∼10% in REV1-depleted extract and ∼25% in Polκ-depleted extract (Figure S5I). This increase in deletion frequency was accompanied by a shift in the distribution of deletions toward longer lengths (Supplementary Figure 5H). Thus, although REV1– and Polκ-dependent TLS past a clb-A adduct introduces frequent point mutations, it appears to suppress an alternative bypass mechanism that is prone to larger deletions. Notably, depletion of REV1 or Polκ-did not affect the distribution of single nucleotide deletions, suggesting that these deletions are not generated during REV1– and Polκ-dependent TLS past a clb-A adduct (Figure 5E). Combined, these sequencing data indicate that a common REV1– and Polκ-dependent TLS mechanism promotes mutagenic bypass of both unhooked colibactin-ICL remnants and colibactin-induced monoadducts.

## Discussion

Despite intense interest in colibactin as a driver of colorectal cancer and IBD, little is known about the repair pathways that enable cells to tolerate colibactin-induced DNA damage. Here, we used plasmids engineered to contain site-specific colibactin-induced ICLs to determine how these lesions are repaired in *Xenopus* egg extract. Our data indicate that colibactin-induced ICL repair is activated upon convergence of replication forks at the lesion and unloading of the replisome (Figure 6A). The FA pathway then mediates nucleolytic incisions that release the cross-link and generate a DSB, which is likely repaired by homologous recombination, and an ICL remnant, which is bypassed by TLS. TLS past the unhooked ICL adduct depends on REV1 and Polκ (most likely in complex with Polζ), which performs both insertion opposite the colibactin-adducted base and extension of the nascent strand. A similar TLS mechanism also enables bypass of colibactin-monoadducts encountered during replication (Figure 6B). Bypass of colibactin-adducts can be error-free but frequently introduces T>A point mutations at alkylated positions. Importantly, we observe no significant differences in the unhooking and bypass mechanisms for the c742-ICL and clb-ICL, validating the use of the c742 analog for investigations into the genotoxic effects of colibactin exposure. Overall, our data provide a comprehensive model for replication-coupled processing of colibactin-induced DNA damage.

**Figure 6.**
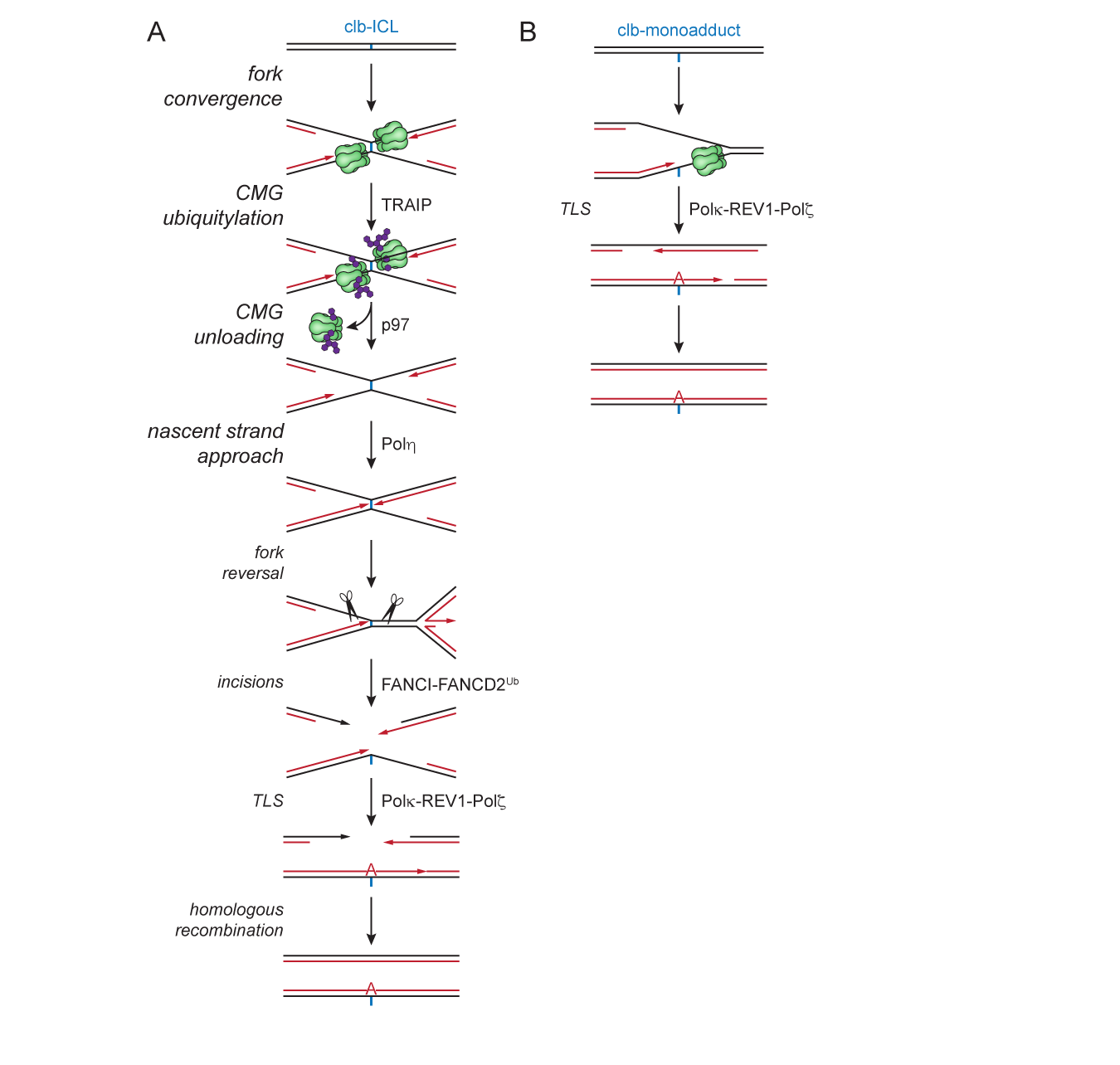
Model for colibactin-induced ICL repair by the FA pathway. (see main text for details) A. Fork convergence at a clb-ICL activates CMG ubiquitylation and unloading. Polη promotes extension of nascent leading strands up to the ICL. Following reversal of one fork, mono-ubiquitylated FANCI-FANCD2 directs nucleolytic incisions that unhook the ICL and generate a DSB. A TLS polymerase complex including REV1-Polκ enable mutagenic extension of the nascent strand past the unhooked ICL remnant, potentially introducing a T>A point mutation at the site of alkylation. The DSB intermediate is then repaired through homologous recombination. B. Replication fork encounters with a colibactin-monoadduct causes DNA unwinding by the CMG helicase to become uncoupled from DNA synthesis by the replicative polymerase. A TLS polymerase complex including REV1-Polκ enable mutagenic extension of the nascent strand past the adduct, potentially introducing a T>A point mutation at the site of alkylation.

### Colibactin-induced ICLs are a physiologically-relevant substrate for the FA pathway

Biallelic mutations in of any 22 *FANC* genes linked to FA cause hypersensitivity to ICL inducing agents^14^. Moreover, the biochemical activities of FANC proteins comprise a common replication-coupled pathway that enables ICL repair. These insights have led to presumption that FA is caused by a failure to repair endogenous ICLs. However, key outstanding question concerns the identities of the cognate endogenous ICLs that require repair by the FA pathway and presumably drive progression of bone marrow failure and cancer in FA patients. Mutations in the FA pathway strongly synergize with mutations in *ALDH2* and *ADH5*, which enable detoxification of acetaldehyde and formaldehyde, respectively^24,25^. These data strongly imply that unrepaired ICLs produced by endogenous reactive aldehydes are causal for FA. However, both the abundance and the specific chemical structures of any aldehyde-induced ICLs that are generated as a consequence of normal human physiology are currently unknown. To our knowledge, the clb-ICL is the only naturally-occurring ICL with strong physical and genetic evidence to substantiate its existence^3,5–9^. Our data therefore demonstrate that the FA pathway does enable repair of an ICL that is likely to arise at some point over the lifetime of most individuals. It is possible, then, that the evolution of the FA pathway has been shaped, at least in part, by exposure to colibactin or another microbiome-derived ICL-inducing agent. Given emerging evidence that the production of genotoxic secondary metabolites may be highly prevalent among bacteria of the human microbiome, it will be interesting to determine whether DNA repair networks are influenced by genetic interactions with microbial genotoxins.

While our data reveal that the FA pathway promotes resistance to colibactin-induced ICLs, it is unclear whether colibactin contributes to the etiology or progression of FA. Given the prevailing view that the uterine environment is generally sterile^42^, it is difficult to envision how colibactin might influence the development of congenital abnormalities associated with FA. Additionally, we are unaware of any evidence linking *pks^+^* bacteria to bone marrow failure. Current data also do not suggest that FA patients are at elevated risk for CRC, though sporadic cases have been reported^43^. Importantly, the colibactin-associated mutation signature has been detected in oral squamous cell carcinomas, a cancer for which FA patients are at dramatically increased risk^44,45^. It will therefore be important to examine the incidence of the colibactin-associated mutation signature in head and neck solid tumors isolated from FA patients. Regardless, our results suggest that FA patients may be at increased risk for complications arising due to colonization by *pks+* bacteria.

### Mechanism of TLS past colibactin-induced DNA adducts

Alkylation by colibactin appears to exhibit a striking specificity for the N3 position of dAs positioned in AT-rich motifs. Although the structure of a colibactin-induced ICL has not been determined, molecular docking suggests that colibactin’s specificity is driven by the toxin’s binding in the relatively narrow minor groove of AT-rich DNA^6^. Alkylation by colibactin ultimately results in the formation of a minor groove adduct. Nucleolytic unhooking of a clb-ICL generates an ICL remnant comprising an N3 alkylated dA linked to a short oligonucleotide through the colibactin scaffold. Adducts with similar structures are expected upon reaction of colibactin with a single dA (monoadduct formation) and upon depurination of a clb-ICL, though the adducts will differ at the cyclopropane distal to the site of N3 dA alkylation. In general, translesion synthesis past minor groove adducts is thought to depend on Polκ. Polκ-deficient cells are hypersensitive to genotoxins (e.g. Illudin S and mitomycin C) that induce minor groove adducts, and Polκ has been shown to efficiently extend DNA primers past *N*^2^-dG adducts (e.g. Benzo[a]Pyrene-*N*^2^-dG) and N3-dA adducts (e.g. 3-deaza-3-methyl-dA) *in vitro*^46,47^. Additionally, TLS past a p3d-Phen-A adduct was recently reported to depend on Polκ polymerase activity in egg extract^41^. In this case, bypass did not require REV1. It is therefore somewhat surprising that the bypass of colibactin-induced adducts requires REV1 and presumably not Polκ polymerase activity. Despite having a catalytic active site that can accommodate minor groove adducts^48^, Polκ might be inhibited by the larger size of the colibactin adducts (clb-A molecular weight ∼900 Da). Colibactin may also make additional contacts to DNA in the minor groove that impede insertion by Polκ. Finally, we note that previous studies have examined bypass of N3-dA adducts using stabilizing 3-deaza-substituted dA analogs that may not completely recapitulate the effects of N3 alkylation.

Our data also indicate that Polη promotes approach of nascent leading strands to within 1 nt of a colibactin-induced ICL. Based on the timing of nascent strand approach product accumulation, we infer that Polη helps to extend the nascent strands after CMG unloading and before FANCI-FANCD2-dependent incisions that unhook the ICL. REV1 and Polκ then act together to enable insertion opposite the alkylated base and subsequent extension past the lesion. Similar coordination between Polη and REV1-Polζ has been observed during bypass of DNA-protein cross-links (DPCs), cisplatin intrastrand cross-links, and UV-induced 6,4 photoproducts^49–51^. In these cases, Polη normally performs insertion of a nucleotide opposite the lesion while REV1-Polζ performs the extension step, though REV1-Polζ can compensate for a Polη deficiency during the insertion step. In the case of colibactin-ICLs, however, Polη appears unable to perform insertion, even when REV1 is absent. It may be then that the structure of the colibactin-ICL blocks insertion by Polη. In the context of model ICL duplexes, strand displacement synthesis and insertion by Polη was shown to be inhibited by a non-distorting ICL, resulting in an accumulation of –1 stall products^52^. Given its long linker length, the colibactin-ICL may also be sufficiently non-distorting to prevent Polη from inserting a nucleotide opposite the cross-linked position and necessitate insertion by Polκ-REV1-Polζ once unhooking has occurred.

### A mutation signature for replication-coupled colibactin-induced ICL repair

Mutation signatures associated with colibactin exposure include predominantly T>C single base substitutions at AT-rich hexanucleotide sequences and single nucleotide T deletions at T-homopolymer sequences^5–7^. T>G and T>A single base substitutions are also detected, though less frequently than T>C mutations. Our data provide some insight as to the origin of these mutation signatures. In the contexts of both ICL repair and monoadduct encounters, bypassing colibactin-alkylated dAs frequently introduces a T>A point mutation through the insertion of dA opposite the lesion. This immediately suggests that the T>A mutations observed in cells, organoids, and colonic crypts arise through Polκ-REV1-Polζ-dependent TLS past colibactin adducts. This subset of colibactin-associated single-base substitutions may therefore represent a mutation signature for clb-ICL repair by the FA pathway.

How then might the more prevalent T>C mutations arise? A trivial explanation is that Polκ-REV1-Polζ-dependent TLS past a colibactin adduct produces different mutations in frogs and humans due to divergence of the TLS polymerase complex between species. Alternatively, T>C mutations might be produced through distinct ICL repair mechanisms or through repair of distinct colibactin-induced DNA lesions. ICLs can be unhooked through replication-independent mechanisms involving mismatch repair (MMR) or transcription-coupled nucleotide excision repair (TC-NER)^40,53^. Both pathways would generate a ssDNA gap at the lesion and require TLS past an ICL remnant in the remaining strand. Interestingly, while Polζ has been reported to be dispensable for replication-independent ICL repair, the catalytic activity of Polκ is required^40^. Colibactin-induced ICLs can decompose into DSBs through depurination and AP site cleavage, requiring DNA end joining. In such instances, the activities of alternative polymerases such as Polλ, Polμ, or Polθ might be required to generate compatible DNA ends for ligation^54^. Thus, the colibactin-associated T>N mutation spectrum could reflect an amalgamation of TLS polymerase activities that act in different contexts.

When ICL unhooking or TLS is inhibited, colibactin-induced DNA lesions appear to be bypassed by a mechanism that tends to introduce larger deletions. We propose that replication forks that undergo persistent stalling at a colibactin-ICL can undergo breakage and collapse into DSBs that are repaired through an error prone end-joining pathway such as Polθ-mediated end-joining (TMEJ)^55^. Similarly, Polθ might enable microhomology-mediated gap skipping at colibactin-induced monoadducts when TLS is inhibited^56^. Consistent with this interpretation, we observe that virtually all of deletions spanning more than 6 base pairs encompass at least a portion of the colibactin alkylation hotspot. Such deletion structures are explainable by Polθ-dependent annealing and extension of 3’ ends at sites of microhomology. It will therefore be interesting to determine whether inhibition of Polθ and other end-joining factors influences frequencies and structures of longer deletions produced during colibactin-induced ICL repair. Going forward, it will be important to examine the mutagenicity of colibactin exposure in cells carrying mutations in *FANC* genes, Polκ, Polζ, Polθ, and other TLS polymerases, and alternative ICL repair pathways. Such experiments will be critical for unambiguously mapping colibactin-associated mutation signatures to specific DNA damage tolerance mechanisms.

## Acknowledgments

We thank J. Campbell and members of the Semlow, Balskus, Herzon, and Duxin labs for comments on the manuscript and P. Knipscheer for technical advice. FANCD2 and REV1 antibodies were a gift from J. Walter. D.R.S. is supported by NIH grant no. R01GM151410 and a Shurl and Kay Curci Foundation Research Grant and is a Ronald and JoAnne Willens Scholar. S.B.H. is supported by NIH grant no. R01CA215553. E.P.B. is supported by NIH grant no. R01CA208834 and is a Howard Hughes Medical Institute Investigator. J.P.D. is supported by Novo Nordisk Foundation Grants no. NNF14CC0001 and no. NNF220C0074140. J.W.H.W. was supported by the A*STAR NSS (PhD) predoctoral fellowship.

## Author contributions

M.V.A. performed the experiments described in Figures 1C–F, 2, as well as Supplementary Figures 1C, D, F, G and 2A, D–F. D.R.S. performed the experiments described in Figures 2A, 3, and 4, as well as Supplementary Figures 1B, E, H; 2B; 3; and 4. M.V.A and X.H. performed and analyzed the experiments described in Figures 5 and Supplementary Figure 5. X.H performed the experiment described in Supplementary Figure 2C. E.J.M., T.Y.W., and T.F.C. assisted with performing and analyzing the experiment described in Figure 2A. M.V.A. performed data analysis for Figure 5 and Supplementary Figure 5, C, D, H, and I. K.M.W. and S.B.H. supplied the c742 analog. J.W.H.W. and E.P.B provided the DNA duplex containing clb-ICL. S.S.B. and J.P.D provided Polη and Polκ antibodies and the pCMV-Sport6-POLH plasmid. M.V.A and D.R.S. designed the experiments. D.R.S. wrote the manuscript with input from all authors.

## Methods

Oligonucleotide sequences used for cross-link generation and sequencing ladder preparation can be found in Supplemental Table 1. Next generation sequencing read counts can be found in Supplemental Tables 2-6.

### Preparation of *Xenopus* egg extracts

Animal work performed at Caltech was approved by the the IACUC (Protocol IA20-1797 approved 28 May 2020). The institution has an approved Animal Welfare Assurance (no. D16-00266) from the NIH Office of Laboratory Animal Welfare. Preparation of high-speed supernatant (HSS) and nucleoplasmic extracts (NPE) from Xenopus laevis eggs was performed as described previously^57^. Briefly, HSS was prepared from eggs collected from six laboratory bred wild-type adult female *X. laevis* (aged >2 years). Eggs were dejellied in 1 L of 2.2% (w/v) cysteine, pH 7.7, washed with 2 L 0.5X Marc’s Modified Ringer’s solution (2.5 mM HEPES-KOH (pH 7.8), 50 mM NaCl, 1 mM KCl, 0.25 mM MgSO_4_, 1.25 mM CaCl_2_, 0.05 mM EDTA) and washed with 1 L of Egg Lysis Buffer (ELB) (10 mM HEPES-KOH (pH 7.7), 50 mM KCl, 2.5 mM MgCl_2_, 250 mM sucrose, 1 mM dithiothreitol (DTT), and 50 μg/mL cycloheximide). Eggs were then packed in 14-mL round-bottom Falcon tubes at 180 x g using a Sorvall ST8 swinging bucket rotor. Eggs were supplemented with 5 μg/mL aprotinin, 5 μg/mL leupeptin, and 2.5 μg/mL cytochalasin B and then crushed by centrifugation at 20,000 x g for 20 min at 4°C in a TH13-6×50 rotor using a Sorvall Lynx 4000 centrifuge. The low-speed supernatant (LSS) was collected by removing the soluble extract layer and supplemented with 50 μg/mL cycloheximide, 1 mM DTT, 10 μg/mL aprotinin, 10 μg/mL leupeptin, and 5 μg/mL cytochalasin B. This extract was then spun in thin-walled ultracentrifuge tubes at 260,000 x g for 90 min at 2°C in a TLS-55 rotor using a tabletop ultracentrifuge. Lipids were aspirated off the top layer, and HSS was harvested, aliquoted, snap frozen in liquid nitrogen and stored at −80 °C. NPE preparation also began with extracting LSS, except eggs were collected from 20 laboratory bred wild-type female *X. laevis* (aged >2 years), and the volumes used to dejelly and wash the eggs were doubled (2 L of 2.2% cysteine, 4 L of 0.5X Marc’s Modified Ringer’s solution, and 2 L of ELB). LSS was supplemented with 50 μg/mL cycloheximide, 1 mM DTT, 10 µg/mL aprotinin, 10 µg/mL leupeptin, 5 µg/mL cytochalasin B, and 3.3 μg/mL nocodazole. The LSS was then spun at 20,000 x g for 15 min at 4 °C in a TH13-6×50 rotor using a Sorvall Lynx 4000 centrifuge. The top, lipid layer was removed, and the cytoplasm was transferred to a 50-mL conical tube. ATP regenerating mix (2 mM ATP, 20 mM phosphocreatine and 5 μg/mL creatine phosphokinase) was added to the extract. Nuclear assembly reactions were initiated by adding demembranated *X. laevis* sperm chromatin^55^ to a final concentration of 6,600 units/µL. After 75–90 min incubation, the nuclear assembly reactions were centrifuged for 3 min at 18,000 x g at 4°C in a TH13-6×50 rotor using a Sorvall Lynx 4000 centrifuge. The top, nuclear layer was then harvested and spun at 260,000 x g for 30 min at 2°C in a TLS-55 rotor using a tabletop ultracentrifuge. Finally, lipids were aspirated off the top layer, and NPE was harvested, aliquoted, snap frozen in liquid nitrogen and stored at −80 °C.

### Protein expression and purification

Recombinant *Xenopus* FANCI-FANCD2 complex was expressed and purified essentially as previously described^36^. The three required recombinant pFastBac1 donor plasmids – the first containing Xenopus FANCI with an N-terminal FLAG-tag, the second containing Xenopus FANCD2 with an N-terminal Strep-tag, and the third containing the ubiquitylation-dead mutant FANCD2^K562R^ with an N-terminal Strep-tag – were generous gifts from the Walter Lab^36^. These donor plasmids were transformed into DH10Bac *E. coli* per the instructions in the Bac-to-Bac Baculovirus Expression System user manual (Invitrogen). After 3 rounds of expression, two in adherent and one in suspension cell cultures, 500 mL Sf9 insect cell suspension cultures were co-infected with viruses expressing either FLAG-xlFANCI and Strep-xlFANCD2 or FLAG-xlFANCI and Strep-xlFANCD2^K562R^ and incubated for 72 hours at 27°C. Cells were collected by spinning at 500 x g for 10 minutes at 4°C and fully resuspended in 1X PBS (137 mM NaCl, 2.7 mM KCl, 10 mM Na_2_HPO_4_, 1.8 mM KH_2_PO_4_). Resuspended cells were spun again at 1,000 x g, supernatant was removed, and cell pellets were frozen in liquid nitrogen and stored at –80°C. To purify the recombinant protein complexes, cell pellets were thawed on ice and resuspended in lysis buffer 10 mL lysis buffer (50 mM Tris-HCL pH 8.0, 100 mM NaCl, 0.5 mM PMSF, 1 mM EDTA, 1 tablet EDTA-free protease inhibitor cocktail (Roche) in 50 mL). The cells were passed through a 21G needle 3 times and lysed by sonication at 40% output for 1 minute (1 second on, 1 second off) for 4 rounds with 20 seconds between rounds. All subsequent steps were performed at 4°C. After centrifugation at 40,000 x g (18,000 rpm) for 40 minutes, the soluble fraction was collected and filtered through a 0.45 µm syringe filter. The filtrate was incubated for 2 hours with 200 µL anti-FLAG M2 affinity resin (Sigma) that was prewashed with lysis buffer.

During the incubation, a 5 mL polypropylene column was packed with 0.5 mL Strep-Tactin XT 4Flow resin (IBA Life Sciences), overlaid with 0.5 mL Strep wash buffer (100 mM Tris-HCl pH 8.0, 150 mM NaCl, 1 mM EDTA), and stored at 4°C until use. After the incubation, the anti-FLAG beads with bound protein complex were washed once with 10 mL lysis buffer and 4 times with 10 mL FLAG wash buffer (20 mM Tris-HCl pH 8.0, 150 mM NaCl, 10 µg/mL aprotinin/leupeptin) by spinning at 1,000 x g for 2 minutes each time. The resin was transferred to 1 mL polypropylene column, and the FANCI-FANCD2 complex was eluted with FLAG wash buffer containing 100 µg/mL 3X FLAG peptide (Sigma) and 5% glycerol. The resin was incubated for 15 minutes in 200 µL elution buffer before each fraction was collected, with the exception of the first fraction, for which the incubation time was 40 minutes. Fractions containing the majority of the protein were pooled and applied to the Strep-Tactin column, which was pre-washed with 1 mL (2 column volumes) of Strep wash buffer. The flow-through was collected and re-applied to the column, which was then incubated for 30 minutes. The column was subsequently washed 5 times with 0.5 mL of Strep wash buffer, and the protein complex was eluted with Strep wash buffer containing 50 mM biotin in 0.5 CV fractions. Fractions containing the majority of the protein were pooled and dialyzed against freezing buffer (20 mM Tris-HCl pH 8.0, 150 mM NaCl, 20% glycerol) overnight. The protein complex was concentrated to ∼5 µM with a 0.5 mL Amicon filter unit (10 kDa MWCO) and stored at –80°C. Where indicated, rFANCI-D2 or rFANCI-D2^K562R^ was added to replication reactions to a final concentration of 375 nM in extracts. Recombinant Polη was prepared using Promega TNT Quick Coupled Transcription/Translation system according to the manufacturer’s instructions. 2 µg pCMV-Sport6-POLH plasmid was mixed with 80 µL TNT master mix supplemented with 20 µM methionine in 100 µL total volume. The reaction mixture was incubated at 30 °C for 90 minutes. The reaction was then concentrated to ∼25 µL by centrifugation at 14,000 x g for 20 minutes in an Amicon 0.5 mL filter unit (30 kDa MWCO). Aliquots were then frozen in liquid nitrogen and stored at –80 °C. 0.25 volumes of the rPolη in vitro transcription/translation mixture were used to supplement 1 volume of NPE.

### Immunodepletions

Immunodepletions using antibodies against FANCD2, REV1 (–N and –C), Polη, and Polκ were performed as previously described^22,36,41,49^. Briefly, Protein A Sepharose Fast Flow (Cytiva) resin was washed with 1X PBS and incubated with an appropriate volume of antibody overnight at 4°C (3.6 volumes of FANCD2, 1 volume of REV1-N and REV1-C, and 5 volumes of Polκ and Polη antibodies). The next day, the beads were washed twice with 1X PBS, once with ELB (2.5 mM MgCl_2_, 50 mM KCl, 10 mM HEPES, 250 mM sucrose), twice with ELB supplemented with 0.5 M NaCl, and 3 further times with ELB. For FANCD2 depletion, 2 rounds of depletion were performed by adding 7.14 volumes of egg extracts (HSS and NPE separately) to 1 volume of beads and incubating on a rotating wheel at room temperature for 20 mins for each round. For REV1 depletion, 5 volumes of egg extracts were added to 1 volume of beads and incubated at 4°C for 60 minutes per round; 2 rounds of depletion were performed with REV1-N beads, followed by one round with REV1-C beads. For Polκ and Polη depletions, 5 volumes of egg extracts were added to 1 volume of beads and incubated at room temperature for 15 minutes per round; one round of depletion was performed for HSS and 3 rounds for NPE. Between rounds of depletion, the samples were spun at 2,500 x g for 30 seconds in an S-24-11-AT rotor in an Eppendorf 5430R, and extract supernatants were collected carefully to prevent bead contamination.

### Preparation of oligonucleotide duplexes with site-specific interstrand cross-links

Site specific cross-links were prepared as previously described^13,58^. To generate the c742-ICL containing oligonucleotide duplex, complementary oligonucleotides (c742-ICL top and c742-ICL bottom; 151 µM each) were mixed in 10 mM NaPO_4_ buffer (pH 6.0). The mixture was heated to 95°C for 5 min and then cooled at 1°C/min to 18°C. Cross-linking was performed by incubating 40 µM annealed duplex with 30 µM c742 colibactin analog in 10 mM NaPO_4_ buffer (pH 6.0) at 37 °C for 4 hours. The mixture was adjusted to 10 mM NaOH, 100 mM NaCl and the cross-linked oligonucleotide duplex was then purified over a Mono Q 5/50 GL column using a gradient from 550 mM to 700 mM NaCl in 10 mM NaOH over 40 column volumes. Fractions containing the cross-linked duplex were combined and buffer exchanged into 10 mM NaOH, 100 mM NaCl with a 4 mL Amicon filter unit (10 kDa MWCO). A second round of purification was performed over a Mono Q 5/50 GL column using a gradient from 550 mM to 700 mM NaCl in 10 mM NaOH over 40 column volumes. Fractions containing the cross-linked duplex were combined and buffer exchanged into 10 mM Tris-HCl (pH 8.8) with a 4 mL Amicon filter unit (10 kDa MWCO). Aliquots were frozen in liquid nitrogen and stored at –20 °C.

To generate the clb-ICL containing oligonucleotide duplex, complementary oligonucleotides (clb-ICL top and clb-ICL bottom; 20 µM) were mixed 1:1 in annealing buffer (10 mM Tris-HCl [pH 8.0], 50 mM NaCl). The mixture was heated to 95°C and then cooled at 1.5°C/min for 50 minutes (to 20°C). Annealed samples were stored at 4°C and warmed to 25°C prior to use. Overnight cultures of *E.coli* BW25113 + BAC-*pks* grown in LB were back-diluted 1:100 into fresh M9-CAS and shaken at 37°C to an OD_600_ of 0.20. Annealed oligonucleotide samples were added directly into the bacteria culture at a dilution factor of 1:40, and the mixture was dispensed into 96-well plates (200 µL per well; Corning). The plates were further incubated at 37°C without shaking for 5 hours. The contents in the 96-well plates were then recombined and spun down at 4,000 x g for 20 mins, and the supernatant was carefully decanted and filtered through a 0.22 µM PES filter (Corning), flash frozen in liquid nitrogen, and lyophilized to completely dryness. Dried samples were reconstituted in Oligo Binding Buffer (Zymo) and purified using the Oligo Clean & Concentrator™ Kit (Zymo) following the manufacturer’s protocol. The cross-linked oligonucleotide duplex was purified over a Mono Q 5/50 GL column using a gradient from 550 mM to 700 mM NaCl in 10 mM NaOH over 40 column volumes.

Fractions containing the cross-linked duplex were combined and buffer exchanged into 10 mM NaOH, 100 mM NaCl with a 4 mL Amicon filter unit (10 kDa MWCO). The cross-linked duplex was further purified on a 10% acrylamide/bis (19:1), 1X Tris-borate-EDTA (TBE), 8 M urea gel. The gel was stained with SYBR Gold, and the cross-linked duplex was visualized with a Blue-Light Transilluminator. The clb-ICL duplex was excised, eluted from crushed gel slices into 10 mM Tris-HCl (pH 8.8), extracted with phenol:chloroform:isoamyl alcohol (25:24:1; pH 8.0), and ethanol precipitated. The cross-link was dissolved in 10 mM Tris-HCl (pH 8.8) and stored at –80°C.

To generate the Pt-ICL containing oligonucleotide duplex, 1 mM cisplatin was converted to activated monoaquamonochloro cisplatin by incubation in 10 mM NaClO_4_ pH 5.2, 0.95 mM AgNO_3_ for 24 hours at 37 °C in the dark. Cisplatin monoadduct was then generated by incubating 0.125 mM Pt-ICL top oligonucleotide in 5.63 mM NaClO_4_ pH 5.2, 0.375 mM monoaquamonochloro cisplatin (in activation mixture) for 12 min at 37 °C. The reaction was quenched by addition of NaCl to 0.1 M. The monoadducted oligonucleotide was then purified over a Mono Q 5/50 GL column using a gradient from 370 mM to 470 mM NaCl in 10 mM NaOH over 40 column volumes. Fractions containing the monoadducted oligonucleotide were pooled and adjusted to 2 mM MgCl_2_. 1.05 molar equivalents of Pt-ICL bottom oligonucleotide were added and buffer exchange was performed with 100 mM NaClO_4_ using an Amicon Ultra-15 3K filter unit at 4 °C. The oligonucleotides in 100 mM NaClO_4_ were allowed to cross-link by incubation at 37 °C for 48 hours. The cross-linked oligonucleotide duplex was then purified on a Mono Q 5/50 GL column using a gradient from 550 mM to 700 mM NaCl in 10 mM NaOH over 40 column volumes. Fractions containing the Pt-ICL duplex were pooled and concentrated using an Amicon Ultra-15 3K filter unit at 4 °C. The Pt-ICL duplex was further purified on a 20% acrylamide/bis (19:1), 1X Tris-borate-EDTA (TBE), 8 M urea gel. The gel was stained with SYBR Gold, and the cross-linked duplex was visualized with a Blue-Light Transilluminator. The Pt-ICL duplex was excised, eluted from crushed gel slices into TE (pH 8.0), extracted with phenol:chloroform:isoamyl alcohol (25:24:1; pH 8.0), and ethanol precipitated. The cross-link was dissolved in 10 mM Tris-HCl (pH 7.4), 10 mM NaClO_4_ and stored at –80 °C.

To generate AP-ICL containing oligonucleotide duplexes, the complementary oligonucleotides AP-ICL top and AP-ICL bottom were annealed in 30 mM HEPES-KOH (pH 7.0), 100 mM NaCl by heating to 95°C for 5 min and cooling at 1°C/min to 18°C. The annealed duplex was then treated with uracil glycosylase (NEB) in 1X UDG buffer (20 mM Tris-HCl, 10 mM DTT, 10 mM EDTA [pH 8.0]) for 120 min at 37°C followed by extraction with phenol:chloroform:isoamyl alcohol (25:24:1; pH 8.0) and ethanol precipitation. The duplex was then dissolved in 50 mM HEPES-KOH (pH 7.0), 100 mM NaCl and incubated at 37°C for 5 to 7 days to allow cross-link formation. Cross-linked DNA was purified on a 20% acrylamide/bis (19:1), 1X TBE, 8 M urea gel. The gel was stained with SYBR Gold, after which the cross-linked products were visualized with a Blue-Light Transilluminator, eluted from crushed gel slices into TE (pH 8.0), extracted with phenol:chloroform:isoamyl alcohol (25:24:1; pH 8.0) and ethanol precipitated. The cross-links were dissolved in 10 mM Tris-HCl (pH 8.5) and stored at –80 °C.

### Preparation of plasmids containing cross-links (pICL)

Plasmids containing ICLs were prepared as described previously^13,15,58^. Briefly, the backbone plasmid (containing 48 lacO repeats) was digested with BbsI in NEBuffer 2.1 for 24 hours at 37°C followed by extraction with phenol:chloroform:isoamyl alcohol (25:24:1; pH 8) and ethanol precipitation. The linearized plasmid was dissolved in TE (pH 8.0) and purified over a HiLoad 16/60 Superdex 200 column using isocratic flow of TE (pH 8.0). Fractions containing the digested plasmid were pooled, ethanol precipitated and dissolved in 10 mM Tris-HCl (pH 8.5). pICL^c742^ was prepared by incubating 0.5 nM BbsI digested plasmid with 1.5 nM c742-ICL duplex and 0.4 U/µL NEB T4 DNA ligase in 1x ligase buffer (50 mM Tris-HCl [pH 8.0], 10 mM MgCl_2_, 1 mM ATP, 10 mM DTT) at 4 °C for 6 hours. The ligation reaction was then concentrated using a 15 mL Amicon filter unit (10 kDa MWCO), extracted with phenol:chloroform:isoamyl alcohol (25:24:1; pH 8), and then buffer exchanged into 10 mM Tris-HCl (pH 8.8), 50 mM NaCl, 1 mM MgCl_2_ with a 4 mL Amicon filter unit (10 kDa MWCO). CsCl was added to the DNA mixture to a homogenous solution density of 1.58 g/mL and ethidium bromide was added to 50 µg/mL. The DNA was then transferred to a Quick-Seal tube and spun for 13 hours at 4 °C in an TLA-100.3 rotor at 279682.5 x g. The covalently closed circular plasmid was collected and extracted with an equal volume of saturated isobutanol to remove ethidium bromide. The DNA was then buffer exchanged into 10 mM Tris-HCl (pH 8.8) with a 4 mL Amicon filter unit (10 kDa MWCO). Aliquots of pICL^c742^ were frozen in liquid nitrogen and stored at –80 °C.

pICL^clb^ was prepared by incubating 0.5 nM BbsI digested plasmid with 0.75 nM clb-ICL duplex and 0.4 U/µL NEB T4 DNA ligase in 1x ligase buffer (50 mM Tris-HCl [pH 8.0], 10 mM MgCl_2_, 1 mM ATP, 10 mM DTT) at 4 °C for 16 hours. The ligation reaction was then concentrated using a 15 mL Amicon filter unit (30 kDa MWCO), extracted twice with phenol:chloroform:isoamyl alcohol (25:24:1; pH 8.0) and once with chloroform, ethanol precipitated, and resuspended in 10 mM Tris-HCl (pH 8.8), 50 mM NaCl, 1 mM MgCl_2_. For every 1 mL of the concentrated DNA mixture, 1 g CsCl was added to the mixture. The DNA was transferred to a 5.1 mL Quick-Seal tube (Beckman Coulter), which was supplemented with ethidium bromide to 50 µg/mL and filled to the top with a CsCl-buffer solution consisting of 1 g of CsCl per 1 mL of 10 mM Tris-HCl (pH 8.8), 50 mM NaCl, 1 mM MgCl_2_ buffer. The mixture was spun for 16 hours at 4 °C in an NVT-90 rotor at 60,000 rpm. The covalently closed circular plasmid was collected and extracted 6 times with an equal volume of saturated isobutanol to remove ethidium bromide. The DNA was then buffer exchanged into 10 mM Tris-HCl (pH 8.8), 50 mM NaCl, 1 mM MgCl_2_ with a 4 mL Amicon filter unit (30 kDa MWCO). Aliquots of pICL^clb^ were frozen in liquid nitrogen and stored at –80 °C. pICL^Pt^ and pICL^AP^ were prepared by incubating 0.5 nM BbsI digested plasmid with 0.75 nM Pt-ICL or AP-ICL duplex and 0.4 U/µL NEB T4 DNA ligase in 1x ligase buffer (50 mM Tris-HCl [pH 7.5], 10 mM MgCl_2_, 1 mM ATP, 10 mM DTT) at 4 °C for 16 hours. The ligation reaction was then concentrated using a 15 mL Amicon filter unit (30 kDa MWCO), extracted twice with phenol:chloroform:isoamyl alcohol (25:24:1; pH 8.0) and once with chloroform, ethanol precipitated, and resuspended in TE. For every 1 mL of the concentrated DNA mixture, 1 g CsCl was added to the mixture. The DNA was transferred to a 5.1 mL Quick-Seal tube (Beckman Coulter), which was supplemented with ethidium bromide to 50 µg/mL and filled to the top with a CsCl-TE solution consisting of 1 g of CsCl per 1 mL of TE. The mixture was spun for 23 hours at 20°C in an NVT-90 rotor at 75,000 rpm. The covalently closed circular plasmid was collected and extracted 6 times with an equal volume of saturated isobutanol to remove ethidium bromide. The DNA was then buffer exchanged into 10 mM Tris-HCl (pH 7.5) for pICL^AP^ and 10 mM Tris-HCl (pH 7.5), 10 mM NaClO_4_ for pICL^Pt^ with a 4 mL Amicon filter unit (30 kDa MWCO). Aliquots of both plasmids were frozen in liquid nitrogen and stored at –80 °C.

### Replication reactions

Plasmid replication reactions in *Xenopus* egg extracts were performed essentially as previously described^59^. Briefly, HSS was thawed and supplemented with 20 mM phosphocreatine, 2 mM ATP, 5 µg/mL creatine phosphokinase, and 3 µg/mL nocodazole. Plasmids were added to the HSS mix to a concentration of 7.5 ng/µL and incubated at room temperature for 30 mins to enable licensing. Where indicated, the licensing mix was supplemented with 167-333 nM 3000 Ci/mmoL [α-^32^P]dCTP and/or 400 nM rGeminin. NPE mix was made with 50% NPE, 20 mM phosphocreatine, 2 mM ATP, 5 µg/mL creatine phosphokinase, and 4 mM DTT in ELB. Where indicated, NPE mix was supplemented with 50 µM PHA-767491 HCl or 300 µM NMS-873.

Replication was initiated by addition of 2 volumes of NPE mix to 1 volume of licensing mix. For analysis on native agarose gels, 1 µL samples of the reaction were removed at indicated time points and quenched by addition to 6 µL of replication stop mix (80 mM Tris-HCl, pH 8.0, 8 mM EDTA, 0.13% phosphoric acid, 10% ficoll, 5% SDS, 0.2% bromophenol blue). After all time points were quenched, samples were digested with 2.5 mg/mL proteinase K for 60 mins at 37°C, and replication intermediates and products were resolved on 0.8% agarose, 1X TBE gels. Gels were transferred to an Invitrogen BrightStar-Plus positively charged nylon membrane by capillary action for 5 hours (or overnight), then dried together with the membrane and visualized by phosphorimaging. Autoradiograms were imaged with a Typhoon FLA 9500 (GE Healthcare) using the FujiFilm FLA 9500 user interface v.1.1 and analyzed using Image Lab v.6.4.0. For Southern blots and nascent strand analysis on sequencing gels, 4 µL samples of the reaction were removed at indicated time points and quenched by addition to 40 µL of clear replication stop mix (50 mM Tris-HCl [pH 8.8], 50 mM NaCl, 1 mM MgCl_2_, 0.5% SDS, 25 mM EDTA). After all time points were quenched, samples were digested with 200 µg/mL RNAse A for 30 mins at 37°C and then with 2 mg/mL proteinase K for 60 mins at 37°C. Samples were adjusted to 200 µL with dilution mix (10 mM Tris-HCl [pH 8.8], 50 mM NaCl, 1 mM MgCl_2_), extracted twice with phenol:chloroform:isoamyl alcohol (25:24:1; pH 8.0) and once with chloroform, and ethanol precipitated. DNA was resuspended in 10 µL of 10 mM Tris-HCl (pH 8.8), 50 mM NaCl, 1 mM MgCl_2_ and stored at –20°C (or –80°C for c742-ICL or clb-ICL plasmids intended for Southern blotting). For samples intended for detection of nucleolytic incisions, 3 µL of recovered DNA were incubated with 5 U HincII in 1X NEBuffer rCutsmart in a total volume of 5 µL at 37°C for 60 mins. For samples analyzed by APE1 digestion, 5 µL of recovered DNA were incubated with 1 U APE1 in 1X NEBuffer 4 in a total volume of 10 µL at 37°C for 60 mins. Samples were subsequently run on 0.8% agarose, 1X TBE replication gels as described above.

### Southern blotting

Southern blotting was performed essentially as previously described^36^. Briefly, samples were replicated as described above, and 3 µL of recovered DNA were incubated with 5 U HincII, 1X NEBuffer rCutsmart in a total volume of 5 µL at 37°C for 30 mins. After the incubation, 5 µL of 50 mM Tris-HCl (pH 8.8), 50 mM NaCl, 10 mM EDTA were added to the reaction. The samples was alkylated with 2 µL of 6X alkaline loading buffer (300 mM NaOH, 6 mM EDTA, 18% ficoll, 0.25% xylene cyanol, 0.15% bromocresol green), and DNA was resolved on a denaturing 0.8% agarose, 50 mM NaOH, 1 mM EDTA gel for 16 hours at 1.25 V/cm. After running was complete, the gel was incubated in 0.25 M HCl for 20 mins and in 0.4 M NaOH, 3 M NaCl for 60 minutes. Downward capillary transfer from the gel onto an Invitrogen BrightStar-Plus positively charged nylon membrane was performed overnight in 0.4 M NaOH, 1.5 M NaCl. The membrane was subsequently washed with 4X SSC buffer (0.6 M NaCl, 60 mM sodium citrate; pH 7.0) for 5 mins, and the DNA was cross-linked to the membrane via UV irradiation in a UVP Hybrilinker Oven. The membrane was then incubated with 15 mL ULTRAhyb Ultrasensitive Hybridization Buffer (Invitrogen) for 1-2 hours at 42°C. During this prehybridization, a radioactive probe was made using 50 ng of linearized pCtrl and the Prime-a-Gene Labeling System (Promega) following the manufacturer’s instructions. The probe was then added to the hybridization tube and incubated with the membrane at 42°C for 18 hours. After hybridization, the membrane was washed twice with 1X SSC, 0.7% SDS at 42°C for 10-20 minutes per wash and twice with 0.1X SSC, 0.7% SDS that was pre-heated to 75°C for 10-20 minutes per wash. The membrane was dried briefly on blotting paper and visualized by phosphorimaging as described above.

### Nascent strand analysis

Nascent strand analysis was performed essentially as previously described^35^. Briefly, samples were replicated as described above, and 3 µL of recovered DNA were incubated with 4 U AflIII and 8 U EcoRI in 50 mM Tris-HCl pH 8, 100 mM NaCl, 10 mM MgCl_2_ in a total volume of 5 µL at 37°C for 3 hours. Reactions were stopped by adding 5 µL of Gel Loading Buffer II (Invitrogen). Primers for sequencing ladders were radiolabeled by incubating 200 nM sequencing primer (5’-CATGTTTTACTAGCCAGATTTTTCCTCCTCTCCTG-3’) with 200 nM [γ-^32^P]ATP and 20 units of T4 PNK (polynucleotide kinase) in 1X T4 PNK reaction buffer in a total volume of 50 µL for 30 minutes at 37°C. The T4 PNK was inactivated by heating at 95°C for 2 minutes, the unincorporated radionucleotides were removed using a Micro Bio-Spin column (Bio-Rad) per manufacturer instructions. Sequencing ladders were made using the Thermo Sequenase Cycle Sequencing Kit: 0.5 pmol of radiolabeled primer were mixed with 175 ng of pCtrl, 2 µL of reaction buffer, and 2 µL of sequenase in a total volume of 17.5 µL to make a master reaction mix. 4 µL of master mix were then mixed with 4 µL of ddATP, ddGTP, ddCTP, and ddTTP termination mixes, and the reactions were incubated at 95°C for 3 minutes, followed by 50 cycles of 95°C for 30 seconds, 60°C for 60 seconds, and 72°C for 90 seconds. 8 µL of Gel Loading Buffer II (Invitrogen) were subsequently added to each reaction, and the sequencing ladders were stored at –20°C until use. All samples were heated at 75°C for 5 minutes, snap-cooled in an ice water bath, and resolved on a 7% acrylamide/bis (19:1), 8 M urea, 0.8X GTG buffer (71 mM Tris, 23 mM taurine, 0.4 mM EDTA) gel. Sequencing gels were dried and visualized by phosphorimaging as described above.

### Plasmid pulldown and sample preparation for mass spectrometry

Plasmid pulldowns for the purpose of mass spectrometry were performed essentially as previously described^60^. Briefly, 6 µL of M-280 Streptavidin Dynabeads (Invitrogen) per time point were washed 3 times with 50 mM Tris-HCl (pH 7.5), 150 mM NaCl, 0.02% Tween-20, and 1 mM EDTA (pH 8.0) and subsequently incubated in the same buffer supplemented with 2 pmol of biotinylated LacI per µL of beads for 40 minutes at room temperature on a rotator wheel. The beads were then washed 4 times with pulldown buffer (2.5 mM MgCl_2_, 50 mM KCl, 10 mM HEPES, 250 mM sucrose, 0.25 mg/mL BSA, 0.02% Tween-20), divided into 40 µL aliquots for time points, and stored on ice. Replication reactions were performed as described above, except extracts were supplemented with 40 µg/mL RNAse A and 10 U/µL RNAse T1 and plasmids were replicated at a concentration of 5 ng/µL in extracts (double the standard 2.5 ng/µL concentration). At indicated time points, 8 µL of reaction mixes were quenched into the 40 µL aliquots of Dynabeads (with 4 replicates for each time point) and immediately incubated on a rotator wheel for 30 minutes at 4°C to enable binding of the plasmid’s *lac* array to the LacI-coated beads. Subsequent wash steps were all performed at 4°C. After the incubation, the beads were washed twice with 2.5 mM MgCl_2_, 50 mM KCl, 10 mM HEPES, 250 mM sucrose, and 0.03% Tween-20 and once with the same solution without Tween-20. For each time point, the beads for one of the replicates were suspended in 10 µL of 1X Laemmli loading buffer and analyzed by immunoblotting, and the beads for the other 3 replicates were suspended in 50 µL of 50 mM HEPES-KOH (pH 8.0), transferred to fresh microtubes, and stored on ice until all samples were ready for the next step. Subsequently, the buffer was aspirated from the beads, and they were resuspended in 50 µL of 8 M urea in 50 mM HEPES-KOH (pH 8.0). Subsequent steps were performed at room temperature unless otherwise indicated. Sample were incubated with 20 mM DTT for 15 minutes. Samples were then supplemented with chloroacetamide to 58 mM and incubated at 1,200 rpm on a Thermomixer for 40 minutes, covered in foil to protect from light. Samples were then incubated with 1.5 µg Lys-C at 1,400 rpm on a Thermomixer for 150 minutes. Following the incubation, the supernatants containing the eluted peptides were separated from the beads on a magnetic rack and transferred to fresh tubes (∼60 µL of supernatant per sample). Samples were adjusted to 200 µL with 50 mM ammonium bicarbonate and incubated with 2.5 µg of trypsin at 30°C overnight. The next day, digestions were terminated by adding 20 µL of 10% TFA (trifluoroacetic acid) and adjusted to 0.5 M NaCl by adding 74 µL of 2 M NaCl. Stage tips were prepared by dispensing two C18 disks into each 200 µL tip with a syringe and spinning the tips in 3D-printed adaptors at 500 x g for 3 mins to pack the disks. All subsequent stage tip spins were also performed at 500 x g for 3 mins. Stage tips were activated by spinning with 100 µL 50% ACN (acetonitrile) and equilibrated by spinning with 100 µL 0.1% TFA. Samples were spun at 8,000 x g for 10 mins to pellet any undigested DNA debris. 85 µL of supernatant were applied to stage tips (and spun) 3 times, leaving ∼40 µL of each sample behind to ensure that any undigested DNA remained undisturbed. Stage tips were washed twice with 100 µL of 0.1% TFA, and peptides were eluted twice with 50 µL of 50% ACN, 0.1% TFA. Samples were then dried by SpeedVac, reconstituted in 10uL of 2% ACN/0.2% FA (formic acid) solution, sonicated for 5 minutes, and centrifuged at 16,000 x g for 5 minutes at room temperature. 4 µL of supernatant from each sample were transferred to autosampler vials and analyzed at the Caltech Proteomics Exploration Laboratory. Samples were analyzed by mass spectrometry using a Q Exactive HF mass spectrometer coupled to Easy-nLC1200.

### Proteomic data processing

Raw data files were searched against the provided *Xenopus* database xenla_10_1_xenbase.fasta using Proteome Discoverer 3.0 software based on the SequestHT algorithm. Oxidation / +15.995 Da (M) and deamidated / +0.984 Da (N) were set as dynamic modifications, and carbamidomethylation / +57.021 Da(C) was set as a fixed modification. The precursor mass tolerance was set to 10 ppm, whereas fragment mass tolerance was set to 0.05 Da. The maximum false peptide discovery rate was specified as 0.01 using the Percolator Node validated by q-value. The R package *limma* was used for normalization and statistical analysis of protein abundances^61^. Abundances were log2 transformed and normalized with quantile normalization. For heatmap visualization, normalized abundances were averaged within biological replicate groups and min-max scaled.

### Immunoblotting

Samples in 1X Laemmli loading buffer (generally representing 1-2 µL of replication reaction or 9 µL of a plasmid pulldown sample) were resolved on 10% or 4-15% acrylamide Mini-PROTEAN or Criterion TGX precast gels (Bio-Rad) and transferred to polyvinyl difluoride (PVDF) membranes (Thermo Fisher). Membranes were blocked with 5% nonfat milk in 1X PBS buffer with Tween 20 (PBST) for 1 hour at room temperature, rinsed 4 times with 1X PBST, and incubated with primary antibodies diluted to an appropriate concentration in 1X PBST at 4°C overnight with shaking. Primary antibody dilutions were as follows: Rabbit polyclonal anti-MCM7 (*Xenopus*) Pocono Rabbit Farm and Laboratory rabbit 456, 1:12,000; Rabbit polyclonal anti-CDC45 (*Xenopus*) Pocono Rabbit Farm and Laboratory rabbit 534, 1:20,000; Rabbit polyclonal anti-H3 (human) Cell Signaling Technology catalog no. 9715S, 1:500; Rabbit polyclonal anti-REV1 (C terminus; *Xenopus*) Pocono Rabbit Farm and Laboratory rabbit 714, 1:5,000; Rabbit polyclonal anti-FANCI (*Xenopus*) Pocono Rabbit Farm and Laboratory rabbit 26864, 1:5,000; and Rabbit polyclonal anti-FANCD2 (*Xenopus*) Pocono Rabbit Farm and Laboratory rabbit 20019, 1:5,000; Rabbit polyclonal anti-Polη New England Peptide rabbit 2991, 1:5,000, and Rabbit polyclonal anti-Polκ New England Peptide rabbit 3924, 1:5000^49^. The next day, the membranes were washed with 1X PBST 3 times for 10 minutes each at room temperature, then incubated for 30 minutes at room temperature with goat anti-rabbit peroxidase-conjugate secondary antibody (IgG, H + L) Jackson ImmunoResearch catalog no. 111-035-003 diluted 1:25,000 in 5% nonfat milk in 1X PBST. After the incubation, the membranes were again washed with 1X PBST 3 times for 10 minutes each at room temperature, then incubated with ProSignal Pico Spray chemiluminescence substrate (Prometheus) for 1-3 minutes at room temperature and imaged using a ChemiDoc Imaging System (Bio-Rad). Contrast was occasionally adjusted to improve visualization of bands. Goat anti-rabbit peroxidase-conjugate secondary antibody (IgG, H + L) was manufacturer validated by antigen-binding assay, western blotting and/or enzyme-linked immunosorbent assay (ELISA). All antibodies against *Xenopus* proteins used in this study were validated by western blotting using *Xenopus* egg extracts.

### Preparation of next-generation sequencing libraries

In preparation for next-generation sequencing (NGS), plasmids were replicated as described above. After extraction and ethanol precipitation, samples were resuspended to a concentration of approximately 10-15 ng/µL. 4 µL of recovered DNA were mixed with 200 nM each of forward and reverse primer (5’-CTCTCCTGACTACTCCCAGTCA-3’ and 5’-GGCGGGACTATGGTTGCT-3’), 200 µM dNTPs, and 2 units of Phusion High-Fidelity DNA Polymerase (NEB) in 1X Phusion HF buffer to a total volume of 100 µL. The region of interest around the ICL was amplified by heating to 98°C for 30 seconds, followed by 18 cycles of 98°C for 10 seconds, 55°C for 30 seconds, and 72°C for 30 seconds, and finished with an incubation at 72°C for 10 minutes. The amplified DNA was purified with a QIAquick PCR Purification Kit (Qiagen) per the manufacturer’s instructions and inspected on a 0.8% agarose, 1X TBE gel visualized with SYBR Gold to confirm amplification. Samples were then submitted to Azenta for next-generation amplicon sequencing. Reads were mapped to a reference sequence, and preliminary analysis was performed by Azenta using NGS Genotyper v1.4.0 software. Depending on the individual data set, 1-12% of mapped reads did not contain any portion of the reference sequence corresponding to the oligonucleotides used to prepare the c742– or clb-ICL inserts. We suspect that these reads were derived from contaminating linear plasmid backbone that underwent end-joiningg (recircularization) during incubation in egg extract. These reads were therefore excluded from further analysis.

## Figure Legends

**Supplementary Figure 1.**
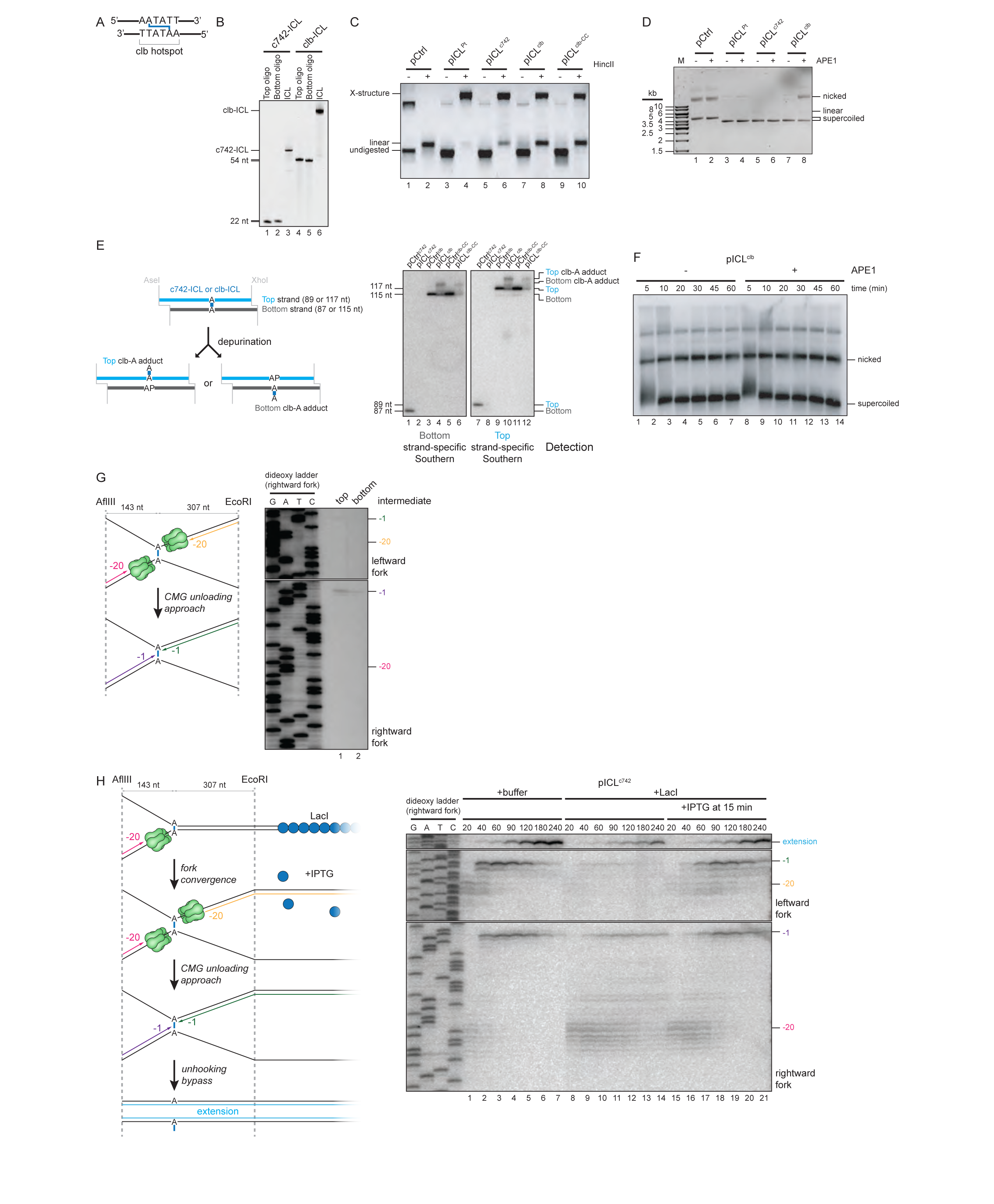
Preparation and replication of plasmids containing a colibactin-induced ICL. A. Schematic of oligonucleotides containing a 5’-AATATT-3’ colibactin alkylation hotspot. The inferred position of the cross-link is indicated. B. Pairs of annealed oligo nucleotides (22 or 54 nt in length) were reacted with c742 or native clb. Cross-linked oligos were FPLC-purified and resolved by denaturing PAGE. C. Plasmids containing the indicated ICLs were linearized with HincII, resolved on a denaturing (alkaline) agarose gel and visualized with SybrGold stain. Note that this method may underestimate the extent of pICL^c742^ and pICL^clb^ cross-linking since it requires extensive incubations, workups, and gel run times, during which the plasmids may undergo depurination. D. Plasmids containing the indicated ICLs were digested with APE1, resolved on a native agarose gel and visualized with SybrGold stain. Note that APE1 cleaves pICL^clb^ (but not pCtrl, pICL^Pt^, or pICL^c742^) indicating the presence of AP sites formed by depurination of the clb-ICL. E. Left, schematic of fragments produced by AseI and XhoI digestion of plasmids containing intact or depurinated colibactin-induced cross-links. Right, the indicated plasmids were digested with AseI and XhoI, resolved on a denaturing PAGE sequencing gel, and visualized by Southern blotting with ^32^P 5’ end radiolabeled strand-specific probes followed by autoradiography. Digestion of uncross-linked control plasmids generates only unmodified top and bottom strands. Digestion of pICL^clb^ plasmids generates a mixture of unmodified and adducted top and bottom strands (since depurination can occur at either cross-linked base). Fragments containing intact cross-links migrate slowly and are not detected in portion of the gel used for blotting. Note that neither unmodified nor modified strands are detected following digestion of pICL^c742^, indicating that the pICL^c742^ cross-link is largely intact. F. pICL^clb^ was incubated in egg extract under non-replicating conditions. The plasmid was recovered at the times indicated, digested with APE1, resolved on a native agrose gel, detected by Southern blotting with ^32^P radiolabeled probes, and visualized by autoradiography. G. Left, schematic of nascent DNA leading strands liberated by AflIII and EcoRI digestion of replication intermediates. Green, CMG helicase. Right, pICL^c742^ replication was replicated in egg extract supplemented with purified top and bottom pICL^c742^ replication intermediates (e.g. Figure 1, C and D) were digested with AflIII and EcoRI and resolved on a denaturing polyacrylamide sequencing gel. Analysis of the nascent DNA indicates that both the rightward and leftward leading strands have stalled one nucleotide upstream of the cross-link (–1 position). H. Left, schematic of nascent DNA leading strand intermediates produced in the presence of LacI and IPTG. Binding of LacI (blue circles) to an array of 48 *lacO* sites adjacent to the ICL blocks progression of the leftward replication fork and prevents convergence at the ICL. Addition of IPTG causes LacI to dissociate and enables progression of the leftward fork up to the ICL. Nascent lagging strands are omitted for clarity. Right, pICL^c742^ was pre-incubated in buffer or LacI and then replicated in egg extract supplemented with [α-^32^P]dCTP. Where indicated, IPTG was added 15 min after initiating replication to dissociate LacI. Replication intermediates were digested with AflIII and EcoRI and nascent DNA leading strands were resolved on a denaturing polyacrylamide sequencing gel. When LacI is bound to pICL^c742^, only the rightward fork encounters and the rightward nascent leading strands stall 20 to 40 nt upstream of the ICL due to the footprint of CMG. Persistent stalling at –20 to –40 is accompanied by a strong reduction in both –1 approach products and full-length extension products, indicating that a single replication fork is insufficient to trigger resolution of the ICL and maturation of the leading strands. Addition of IPTG at enables the leftward fork to reach the ICL. Arrival of the leftward fork triggers approach of the nascent leading strands up to the ICL and subsequent formation of full-length extension products. This result indicates that pICL^c742^ unhooking is activated by convergence of two replication forks on either side of the ICL.

**Supplementary Figure 2.**
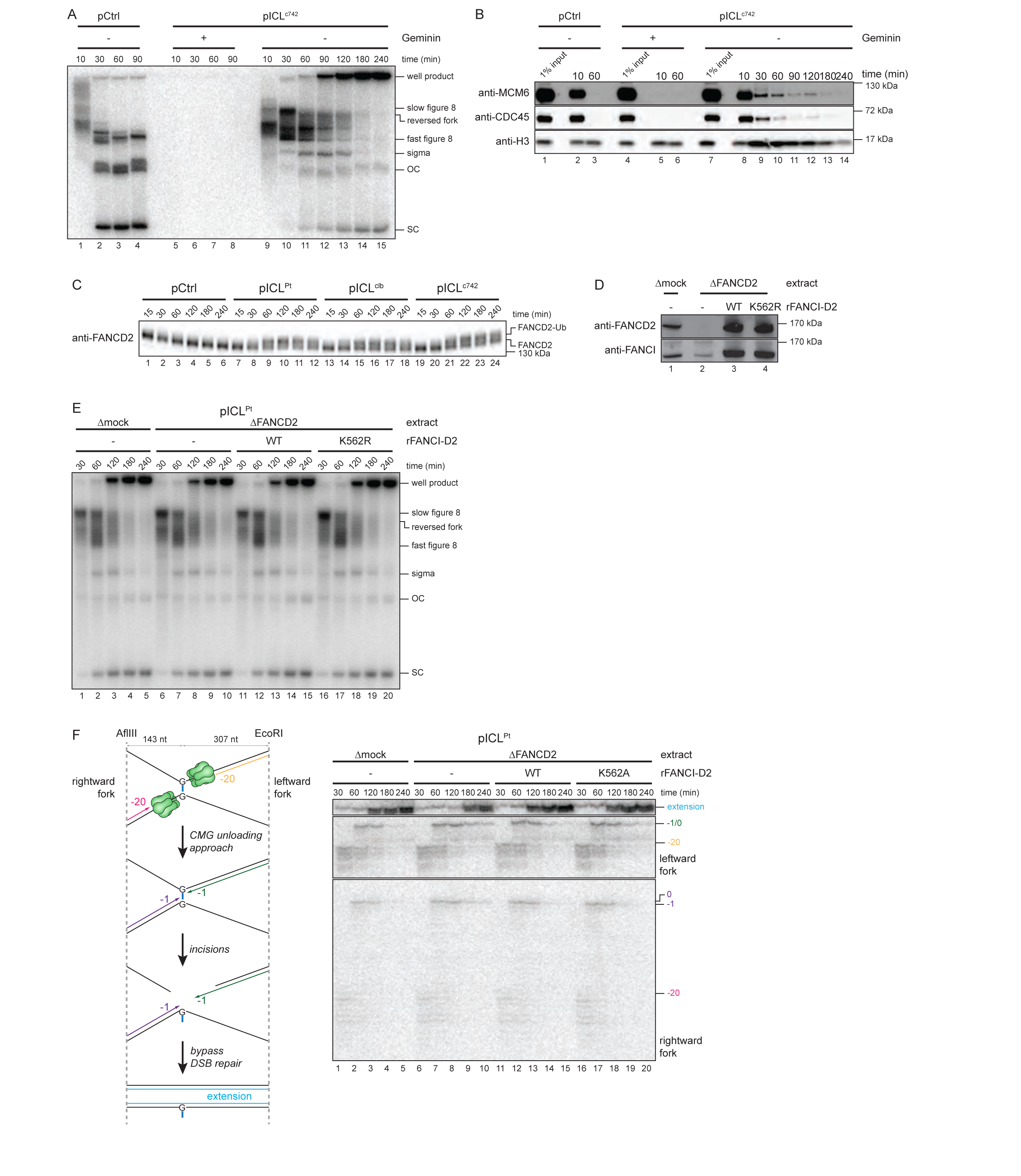
Colibactin-induced ICLs mobilize the FA pathway. A. In parallel with the reactions analyzed in Figure 2A, reactions were supplemented [α-^32^P]dCTP and the replication intermediates were analyzed as in Figure 1D. SC, supercoiled; OC, open circular. B. In parallel with the reactions analyzed in Figure 2A, chromatin-associated proteins recovered from extract were separated by SDS-PAGE and the indicated proteins were detected by immunoblotting. C. The indicated plasmids were replicated in egg extract and, at the times indicated, extract proteins were separated by SDS polyacrylamide gel electrophoresis and immunoblotted for FANCD2. In the presence of an ICL, FANCD2 becomes monoubiquitylated, resolvable as a slower migrating protein band. D. FANCI-FANCD2 immunodepletion. The extracts used in the replication reactions shown in E and F were blotted for FANCI and FANCD2. E. pICL^Pt^ was replicated with [α-^32^P]dCTP in mock– or FANCI-FANCD2-depleted extracts supplemented with rFANCI-FANCD2, as indicated. Replication intermediates were analyzed as in Figure 1D. SC, supercoiled; OC, open circular. F. Nascent DNA strands from the pICL^Pt^ replication reactions shown in D were isolated at the indicated times, digested with AflIII and EcoRI, and analyzed as in Figure 1F.

**Supplementary Figure 3.**
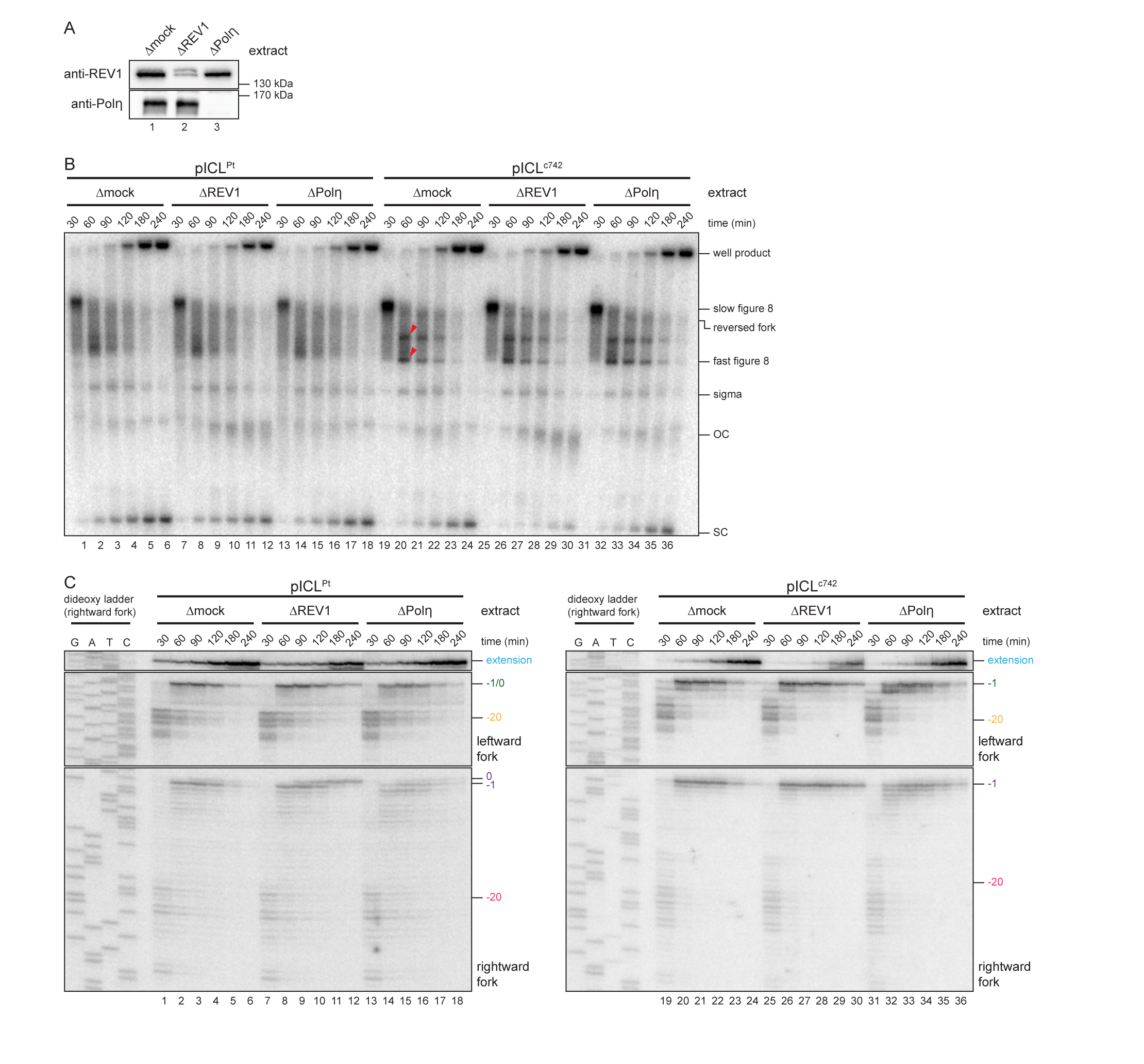
Polη promotes bypass of Pt– and c742-ICLs. A. REV1 and Polη immunodepletion. The extracts used in the replication reactions shown in Supplementary Figure 3, B and C were blotted for REV1 and Polη. B. pICL^Pt^ and pICL^c742^ were replicated with [α-^32^P]dCTP in mock-, REV1-, or Polη-depleted extracts and replication intermediates were analyzed as in Figure 1D. SC, supercoiled; OC, open circular. Red arrowheads indicate top and bottom intermediates formed during replication of pICL^c742^. C. nascent DNA strands from the pICL^Pt^ and pICL^c742^ replication reactions shown in Supplementary Figure 3B were isolated at the indicated times, digested with AflIII and EcoRI, and analyzed as in Figure 1F.

**Supplementary Figure 4.**
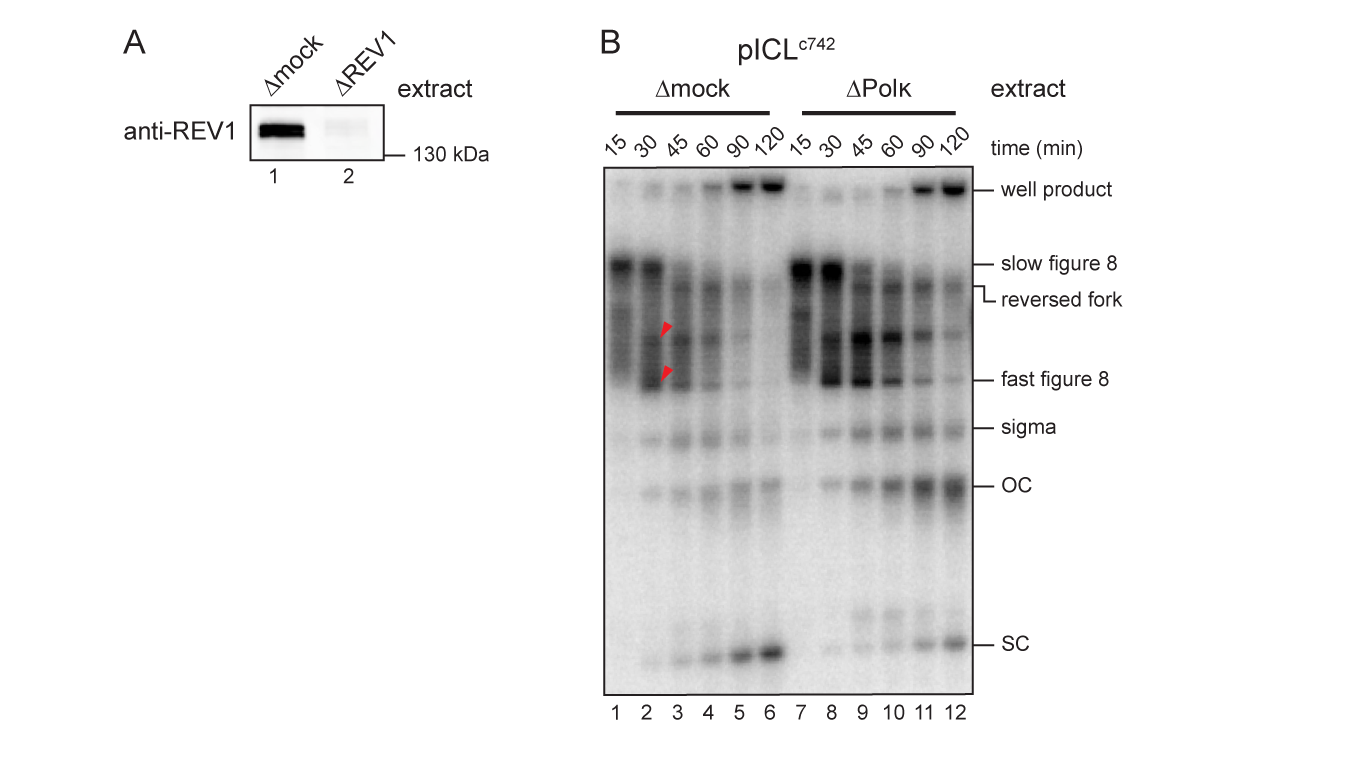
Polκ promotes translesion synthesis past a colibactin-induced monoadduct. A. REV1 immunodepletion. The extracts used in the replication reactions shown in Figure 4, D and E were blotted for REV1. B. pICL^c742^ was replicated with [α-^32^P]dCTP in mock– or Polη-depleted extracts (without p97 inhibitor) and replication intermediates were analyzed as in Figure 1D. SC, supercoiled; OC, open circular. Red arrowheads indicate top and bottom intermediates formed during replication of pICL^c742^. C. pICL^clb^ was replicated with [α-^32^P]dCTP in mock– or Polκ-depleted extracts that were supplemented with p97 inhibitor. Nascent DNA strands were isolated at the indicated times, digested with AflIII and EcoRI, and analyzed as in Figure 1F.

**Supplementary Figure 5.**
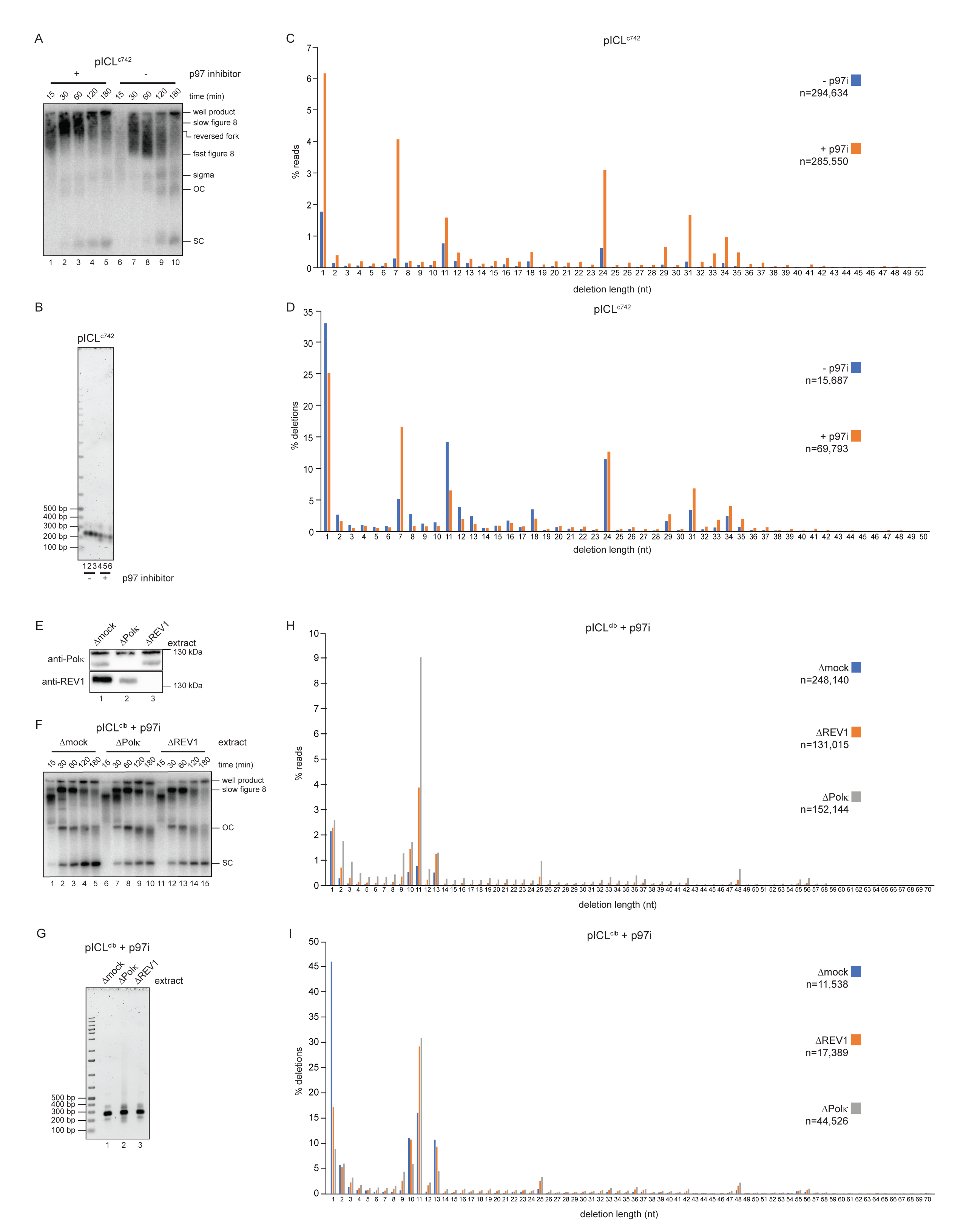
Preparation and of next generation sequencing libraries. A. In parallel with the reactions used to generate sequencing libraries described in Figure 5, B and C and Supplementary Figure 5, B-D, pICL^c742^ was replicated in extract supplemented with [α-^32^P]dCTP and p97 inhibitor, as indicated, and replication intermediates were analyzed as in Figure 1D. SC, supercoiled; OC, open circular. B. In parallel with the reactions shown in Supplementary Figure 5A, pICL^c742^ was replicated for 300 min in the indicated extracts supplemented with p97 inhibitor but lacking [α-^32^P]dCTP. The region of the replicated plasmids surrounding the ICL was then amplified by PCR. PCR amplicons were resolved by native agarose gel electrophoresis and visualized by Sybr Gold staining. The expected length of the PCR amplicons is 240 bp. PCR reactions were performed in triplicate and pooled for amplicon sequencing. C. pICL^c742^ was replicated in extract supplemented with p97 inhibitor, as indicated, and repair products were PCR amplified and sequenced as in Figure 5, B and C. The fraction of sequencing reads corresponding to a deletion of a given length is plotted for each extract condition. n indicates the number of mapped reads obtained for each condition. D. pICL^c742^ was replicated in extract supplemented with p97 inhibitor, as indicated, and repair products were PCR amplified and sequenced as in Figure 5, B and C. The fraction of deletions corresponding to given length is plotted for each extract condition. n indicates the number of mapped deletions observed for each condition. E. REV1 and Polκ immunodepletion. The extracts used in the replication reactions analyzed in Figure 5, D and E and Supplementary Figure 5, F to I, were blotted for REV1 and Polκ. F. In parallel with the reactions used to generate sequencing libraries described in Figure 5, D and E, and Supplementary Figure 5, G to I, pICL^clb^ was replicated in the indicated extracts supplemented with [α-^32^P]dCTP and p97 inhibitor. Replication intermediates were analyzed as in Figure 1D. SC, supercoiled; OC, open circular. G. In parallel with the reactions shown in Supplementary Figure 5F, pICL^clb^ was replicated for 180 min in the indicated extracts supplemented with p97 inhibitor but lacking [α-^32^P]dCTP. The region of the replicated plasmids surrounding the ICL was then amplified by PCR and the amplicons were analyzed as in Supplementary Figure 5B. The expected length of the PCR amplicons is 268 bp. H. pICL^clb^ was replicated in the indicated extract supplemented with p97 inhibitor and repair products were PCR amplified and sequenced, as in Figure 5D. The fraction of sequencing reads corresponding to a deletion of a given length is plotted for each extract condition. n indicates the number of mapped reads obtained for each condition. I. pICL^clb^ was replicated in the indicated extract supplemented with p97 inhibitor and repair products were PCR amplified and sequenced, as in Figure 5D. The fraction of deletions corresponding to given length is plotted for each extract condition. n indicates the number of mapped deletions observed for each condition.

